# TRACKING THE COMPOSITION AND STABILITY OF MICROBIOME ACROSS INDIAN SOCIAL HONEYBEES FORAGING IN A HOMOGENOUS RESOURCE LANDSCAPE

**DOI:** 10.64898/2026.04.19.719467

**Authors:** Dipendra Nath Basu, Pawan Khangar, Kunjan Joshi, Shivani Krishna, Imroze Khan

## Abstract

Microbial communities are essential for host health and ecosystem stability. However, whether host identity or shared foraging resources shapes microbiome structure among co-occurring species remains poorly understood. We studied bacterial and fungal communities of four Indian honeybee species in a mustard monoculture resource condition, integrating behavioural observation-based pollinator data with microbial co-occurrence networks derived from metabarcoding. Microbiome composition was linked to host identity rather than foraging behaviour, bee abundance, or landscape use. While core bacterial taxa were shared, relationships among bacterial cobionts, unlike those among fungal genera, remained species-specific. Microbial diversity, along with community structure and function, influenced network stability, with a highly modular microbial network of *Apis cerana* exhibiting more resilience to simulated perturbations. In summary, host-specific filtering shaped the microbiome more than resource homogenisation, with closely related species facing unique risks of microbial collapse, with broader implications for vulnerability to microbiome imbalance, environmental stress, and emerging infections.

## INTRODUCTION

The functional stability of ecological communities is increasingly threatened by anthropogenic disturbances and climate change, which drive global biodiversity loss and habitat simplification^1,2^. A community’s ability to withstand such stressors depends fundamentally on the structural integrity of its broader ecological networks, including plant-pollinator interactions. The interacting species within these networks do not operate in isolation; rather, they function as biological hosts to complex microbial assemblages. Because these linkages span multiple trophic levels, environmental perturbations can trigger cascading effects across ecological assemblages—disrupting external host-host interactions, altering host-microbe associations, and ultimately destabilising internal microbe-microbe dynamics^3,4^. At this foundational level, microbial communities serve as mediators that enhance host resilience, facilitate nutrient cycling, and stabilise the interspecific relationships of their hosts^5–7^. Consequently, disruption of a host’s microbial community can reduce the fitness of the conspecific population and further weaken the overarching ecological networks to which it contributes^8–10^. Although microbial sharing among individuals and populations is recognised as a factor shaping health and environmental adaptability^11–13^, the structural properties of these microbial metacommunities and their vulnerability to disturbance remain insufficiently characterised, especially within complex social organisms inhabiting shared environments.

Social insects, particularly pollinators such as honeybees, provide an effective model system for investigating such microbial dynamics. Bees are essential for the maintenance of over 70% of known crops and global plant networks^14^. Their ecological interactions occur at multiple, overlapping levels: within colonies, between colonies, and interspecifically at shared floral resources. Flowers function as key environmental interfaces where multi-species foraging promotes extensive microbial exchange^15^. As a result, the microbiomes of social bees may not be assembled in isolation but can be shaped by both host physiology and the broader foraging environment. Disturbances to these pollinator microbiomes, often resulting from agricultural chemicals, habitat homogenization/loss, and climate change, can reduce physiological functions, including digestion, immune defence, and detoxification, ultimately affecting individual fitness and threatening the persistence of essential pollination services^16,17^.

Despite the broader significance of bee-microbe interactions, substantial gaps persist in our understanding of microbiome community assembly and network stability in honey bees. Existing meta-analyses of recent studies demonstrate that European and American eusocial bee species possess a highly conserved core bacterial gut microbiota (i.e., common bacterial taxa shared across all bee species)^18,19^. However, non-random variation in broader microbial networks in paleotropical honeybees, due to (1) transient environmental acquisition of microbiota by hosts, (2) changing interspecific interactions among hosts in response to environmental variation, and (3) interactions with largely uncharacterized fungal communities, remains unexplored. There is a lack of comparative insight into whether closely related, co-occurring species maintain distinct microbiome compositions or exhibit greater microbial sharing while foraging in highly homogenised environments, with broader implications for host and pollinator community health. Additionally, most pollinator microbiome studies focus on bees from temperate regions, such as North American and European social honey bees, bumble bees, and managed apiary species ^20–22^, resulting in limited comparative knowledge of paleotropical communities that support extensive native bee diversity that are ecologically distinct in terms of climatic regimes, seasonal phenology, and dependency on floral resources. It is not yet clear whether shared environmental exposure in these altered resource conditions and landscape heterogeneity supersedes host-specific filtering, or how the resulting microbial co-occurrence networks influence a community’s structural resilience to dysbiosis (i.e., a microbial imbalance within or on the body resulting from a reduction in beneficial microorganisms or overall decrease in microbial diversity).

To address these knowledge gaps, we characterised the internal bacterial and fungal communities of four co-occurring social honey bee species (*Apis cerana, A. dorsata, A. florea*, and *A. mellifera*) foraging within a semi-arid, seasonal monocultural agroecosystem in northwestern India. In this region, mustard is a mass crop, and during the peak bloom in winter, it becomes the sole agricultural floral resource, where foraging resources temporarily converge for most insect species. Presence of a single flowering resource drives a pollinator network towards the highest connectance and lowest modularity, thereby increasing hosts’ ability to share microbes in a natural environment^23^. We asked whether honeybee microbiome assembly is driven more by shared foraging environments or by host-specific microbial filtering. Three interconnected hypotheses were tested regarding the structural assembly and stability of these metacommunities: (i) whether floral resource homogeneity and associated foraging behaviour of honey bees influence their microbiome profiles more than host-specific identities; (ii) if microbiome assembly is influenced by shared resources/resource homogenization, microbial communities should converge across host species and exhibit low modularity. Conversely, if host-specific filtering is more important, microbiomes would exhibit greater compartmentalisation, higher modularity, and stronger within-species than between-species associations. Thus, here we investigate whether networks of bacterial microbiome and fungal cobionts shared similar co-occurrence patterns between honey bee species; (iii) whether these structural shifts in microbial networks directly influence stability of the microbial community under simulated perturbation. By integrating behavioural observations, amplicon-based metagenomics, and Bayesian network inference, this research establishes a comparative framework to understand how host-specific factors influence metacommunity stability in pollinator networks (see the summary of the study design in Fig. 1).

**Figure 1.**
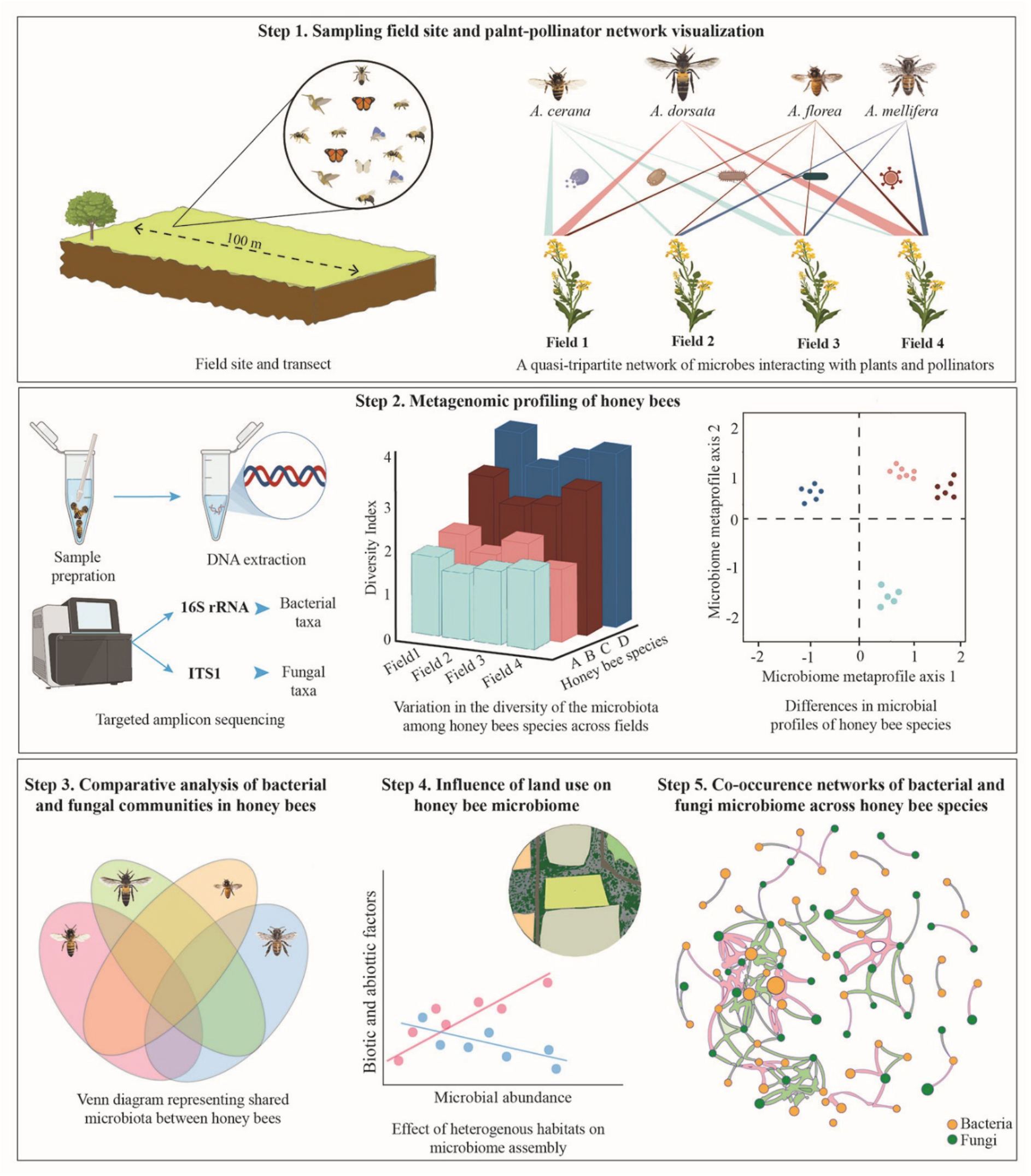
A graphical summary of characterising the microbiome in Indian social honey bees from a homogeneous landscape. **(A)** Sampling of the four common Indian honeybee species and the plant-pollinator network in a homogeneous habitat at Aravalli Hills in northwestern India. **(B)** Characterizing microbiome profile in honey bee species, **(C)** Extent of co-occurrence in microbial taxa within each honey bee species and shared microbiome among the common social honey bee species, **(D)** Correlating behavioural and spatial variability with variation in microbiome profile of Indian social honey bees, **(E)** Predicting modularity and stability of microbial networks.

## RESULTS

### Differences in abundance and foraging among honeybee species suggest heterogeneity in the pollinator visitation pattern

Across the seven study sites (MUST1-7, Fig. S1), we sampled visitation by 3882 insects, mainly from the order Hymenoptera, with a few visitors from Diptera and Lepidoptera. The species diversity (Shannon’s diversity) of flower visitors varied between 1.86 and 2.3 across the different sites, with a high Shannon’s index suggesting equal abundance of all four honey bee species. *A. mellifera* constituted 41% of the honeybees visiting the mustard fields, followed by *A. dorsata* (25.4%), *A. florea* (19.3%), and *A. cerana* (14.11%) (Dataset S1). The abundance (number of bees visiting flowers) and the time spent per flower of the four honeybee species varied significantly across fields (Fig 2A –B; Table S1-S2; in case of both abundance and time spent per flower, Species × Field interaction p-values were <0.001). A species-wise post-hoc comparisons of abundance suggest that these differences were found between all the honeybee species with the exception of *A. florea* and *A. dorsata* (*A. dorsata* – *A. florea*, p = 0.17 (across all the field sites), all other pairwise comparisons were significant: p < 0.05, Table S1). We also found that the abundance of honey bees in field 2 (MUST2) was significantly higher than in the other fields (Table S1). Our results also showed that although the abundance of *A. cerana* is relatively lower than that of *A. dorsata* and *A. mellifera, A. cerana* exhibits greater relative evenness in abundance across fields (Dataset S1).

**Figure 2.**
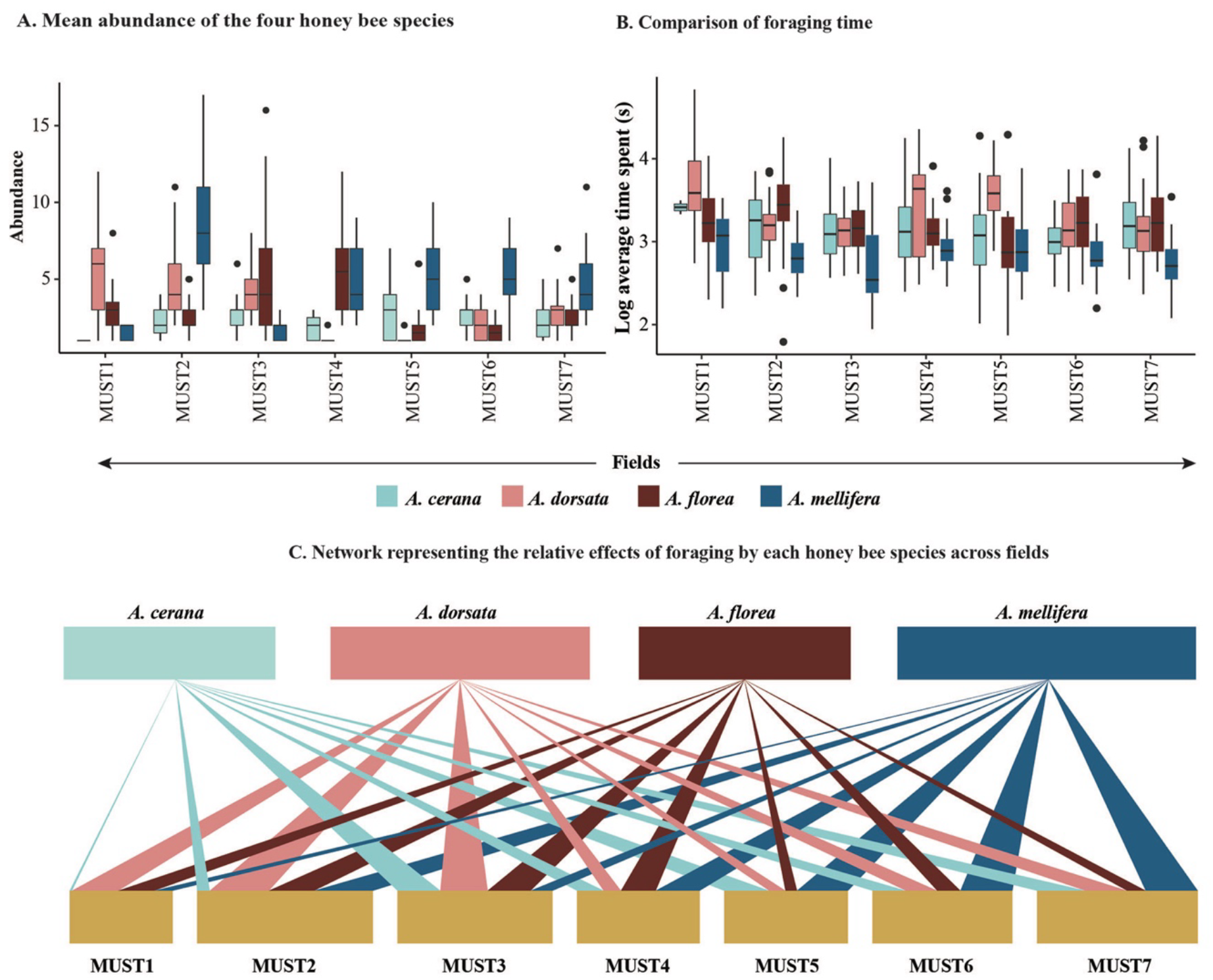
Description of the plant-pollinator network. **(A)** Abundance of the four honey bee species in the seven survey fields, **(B)** A comparison of foraging time (denoted as time spent per flower in each of the fields), **(C)** A summary network representing the relative effects of abundance and foraging by each honey species across the seven fields. Here, the width of the sites (MUST1, MUST2) represents the total abundance of all four honey bee species and the width of the connecting lines denotes abundance of each species in each field.

Similarly, pairwise post hoc comparisons of time spent per flower suggest there are species-wise differences except for *A. florea* and *A. dorsata* (*A. dorsata* – *A. florea*, p = 0.27 (across all the field sites), all other pairwise comparisons: p < 0.05, Table S2). Moreover, foraging time was different between field 1 (MUST1) vs all other fields and field 3 (MUST3) vs field 6 (MUST6) (Dataset S1). Similar to abundance, we found *A. cerana* shows relative evenness in time spent per flower, as we did not find any significant difference across field sites (Table S2). The relative abundance and foraging parameters show the heterogeneity in the relative participation of each honey species in this pollinator network across the seven fields (Fig 2C). Overall, our results suggest that, despite a monoculture resource condition, the four co-occurring honey bee species exhibited significant spatial heterogeneity in their relative abundance and foraging behaviours, reflecting distinct, species-specific engagement within the shared resource space.

### Alpha diversity estimates, comparisons of bacterial and fungal cobionts, and their functional profiles show differential microbial sharing among honey bee species

We performed amplicon sequencing to identify bacterial and fungal cobionts and obtained 27.5 – 197 thousand reads for bacteria and 3.3–50.5 thousand reads for fungi across samples (Dataset S2). Mapping these reads against relevant databases (see methods), we identified 1208 bacterial ASVs, of which 89% were assigned to a unique taxon at the genus level. Similarly, for fungi, we obtained 531 ASVs, out of which 57.4% were assigned genus identification (note that, although lower-order taxon assigning was limited, we obtained a 94% taxon-level identification at the Phylum level). Subsequently, we found the richness of bacterial genera varied significantly between the honeybee species, with a lowest of 13 genera in *A. mellifera* at field 5 and a highest in *A. cerana* with 101 genera at field 7 (Negative binomial model: Richness of genera ∼ Bee species; χ^2^ = 63.417, df = 3, p < 0.001; Fig 3A, Table S3, Dataset S3). However, the fungal richness was homogeneous across the four bee species (Negative binomial model: Richness of genera ∼ Bee species; χ^2^ = 2.759, df = 3, p = 0.43; Fig 3B, Table S3, Dataset S3), with *A. florea* and *A. mellifera* harbouring the least (21 genera) and highest (30 genera) fungal diversity (Fig 3B, Dataset S3), respectively. Moreover, *A. cerana* and *A. dorsta* exhibited lesser variation in their bacterial richness across fields, whereas *A. florea* showed maximum variation in bacterial richness across field sites (min-max scaled fold-change range from 1.6 in *A. cerana* to 3.1 in *A. florea*; Fig 3C, Dataset S3). However, fungal diversity was homogeneous among the four honey bee species (Fig. 3C, Dataset S3).

**Figure 3.**
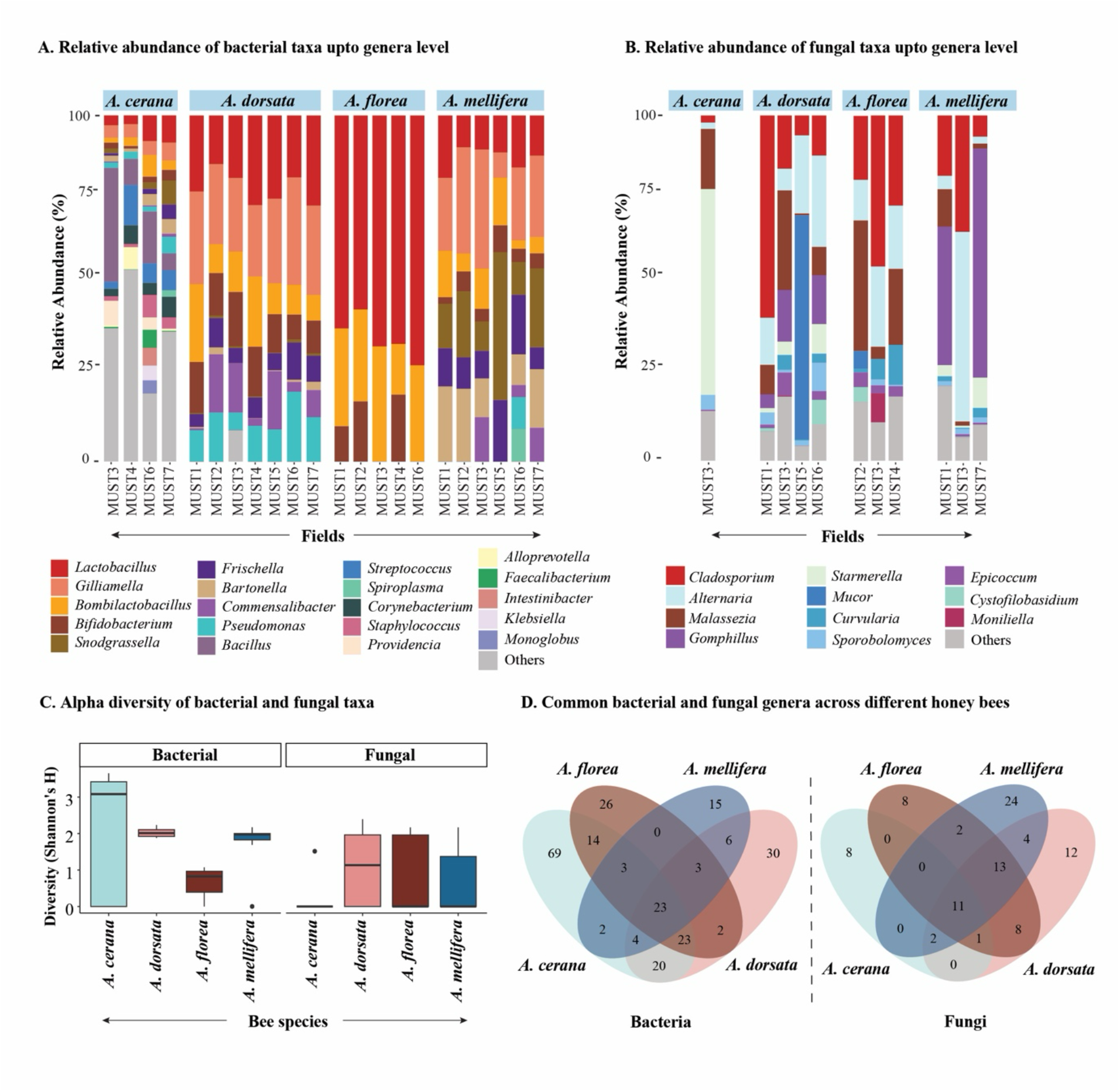
A comparative microbiome profile of the four honey bees. **(A-B)** Relative abundance of the top 20 bacterial and fungal genera from each honey bee species. Microbial abundances vary across fields due to low abundance or absence of honeybee species in that field site, **(C)** Diversity indices of bacteria and fungi across four social honey bee species, **(D-E)** Sharing of bacterial and fungal microbiome among four honey bee species.

We found 38 genera of bacteria, including five bacterial genera consisting of *Gilliamella, Snodgrassella, Bifidobacterium, Lactobacillus* Firm-4, and *Lactobacillus* Firm-5, form the core microbiota, which are found across all four honeybee species at varying abundance (Fig 3D). We also found that *A. cerana* harbours more unique bacterial cobionts and also shares most bacterial genera with *A. dorsata*. In the case of fungi, 11 genera were shared across all four honeybee species, including *Cladosporium, Mucor, Cystofilobasidium*, and *Starmerella* (Fig 3D). We also found *A. mellifera* hosts more unique fungal genera, and fungal sharing is highest between *A. dorsata* and *A. florea*. While examining homogeneity, we found that the bacterial cobiont profile of *A. cerana* is more dispersed, whereas that of *A. dorsata* overlaps with both *A. florea* and *A. mellifera* (Fig 3D). However, for fungal cobionts, we did not find such species-wise dispersions (Fig. 3D). Our results suggest that, despite sharing conserved core taxa, the broader microbial architectures of the four honey bee species diverged significantly, with bacterial communities exhibiting strong host-specific heterogeneity, whereas fungal associations were less distinct.

Subsequently, we also predicted the functional potential of the bacterial communities across the four honeybee species based on the abundances of functional pathways derived from the ASVs using PICRUSt2 software^24^. These pathways provide insights into the metabolic capabilities of microbiome. We subsequently used a canonical variate analysis to obtain differences in the metabolic profile of the microbiome across species. We found that the functional metaprofile of microbes in *A. cerana* is starkly different from those of the other three honey bee species along canonical variate axis 1 (which explains 74.57% of the total variance) driven by functional pathways including N-glycan sysnthesis, Tetracycline synthesis, secondary metabolite synthesis pathways such as isoflavonoid biosynthesis, lipid metabolism pathways such as sphingolipid metabolism and alpha-linolenic acid metabolism and geranoil degradation pathway, intermediates of which are known to affect bacterial quorum sensing (Fig S2A, Table S4, Dataset S4). Moreover, the functional metaprofile of bacterial microbes in *A. florea* differs significantly from those of *A. dorsata* and *A. mellifera* along canonical variate axis 2 (which explains 20.8% of the total variance) driven by pathways related to insulin signalling, ascorbate and aldarate metabolism, dioxin and xylene degradation (Fig S2B, Table S4, Dataset S4).

### Bacterial diversity is independent of the abundance, foraging behaviour of honeybees and anthropogenic landscape heterogeneity

To understand whether variation in cobionts’ numbers is driven by honey bee foraging preferences, we examined the association between microbial abundance and honey bee foraging parameters, such as abundance and time spent on flowers. By and large, we found that the richness of bacterial and fungal genera of the honeybees was not influenced by the foraging parameters, except that time spent per flower was negatively correlated with fungal richness (Fungal richness – time spent: R^2^= 0.471, p-value= 0.02; Fig S3). We further examined landscape use around each sampling field site. The QGIS mapping estimating landscape heterogeneity around field sites show the surroundings of fields 1 and 3 are relatively more heterogeneous compared to the other five field sites, and three of the sites are in close proximity to the forest (Fig S4, Table S5). Subsequently, we estimate the covariations between landscape use and the cobiont abundance using a two-block partial least square method and found that the abundance of both bacterial and fungal cobionts are independent of landscape use heterogeneity (bacterial: RV-coefficient: 0.296 p-value: 0.368; fungal: RV-coefficient: 0.205 p-value: 0.653). These results suggest that microbial richness is not primarily driven by external environmental determinants, functioning largely independent of landscape heterogeneity and foraging behaviour aside from minor, species-specific covariations in fungal microbiome.

### Structural characterisation of both bacterial and fungal communities shows phylogenetically independent non-random co-occurrence patterns of microorganisms

While estimating co-occurrence between bacterial metacommunity including all the four honeybee species, we found 225 bacterial genera co-occurred at varied extents (Fig S5). Species-wise comparison of bacterial co-occurrence showed similar patterns as bacterial richness, with *A. cerana* harbouring more co-occurring bacteria (n=139) and *A. mellifera* hosting the least co-occurring bacterial cobionts (n=43) (Fig S6, Dataset S5). Most of these bacteria show positive co-occurrence across honeybee species, except for a few bacterial genera, such as *Bombilactobacillus, Haemophilus, Acinetobacter*, and *Brevibacterium*, which exhibit only negative associations (Fig S5). Subsequently, we found that bacterial cobionts co-occur independently of their phylogenetic relations across all honey bee species (Fig S7). In the case of fungal cobionts, we found only 73 fungal genera co-occurring across all the honeybee species (Fig S8). A species-wise comparison of fungal co-occurrence showed that *A. mellifera* harbours the most co-occurring fungal cobionts (n=42), whereas *A. florea* hosts the least co-occurring fungal genera (n=25) (Fig S9, Dataset S6). As with bacteria, most fungal genera also show positive co-occurrence across four honeybee species, suggesting these bacterial and fungal genera could be in an associative relationship, potentially associated with environmental or general physiological factors in honey bees.

Subsequently, to obtain community structure, we estimated the co-occurrence networks of the bacterial and fungal metacommunities separately. We found 10 modules of the bacterial community, seven of which are interconnected by core bacterial genera, including *Lactobacillus, Copromonas, Bartonella, Frischella, Gilliamelia, Anaerococcus, Lawsonella, and Fusobacterium* (Fig S10A). The other three modules, comprising 14 bacterial genera, are isolated from other modules but with positive intramodular co-occurrence. While comparing bacterial co-occurrence networks across honeybee species, we found that the co-occurrence network in *A. cerana* exhibits the highest modularity and the lowest nestedness, suggesting that the community is organised into distinct, specialised, and functionally independent sub-communities, leading to niche partitioning and localised interactions. However, in *A. florea* the bacterial network is least modular with the highest connectance, and in *A. mellifera* the network shows the highest nestedness (Fig 4, Table S6). Moreover, the bacterial cobionts exhibit very few shared co-occurrences across four honey bee species, suggesting more species-specific microbial community structures (Fig S10A).

**Figure 4.**
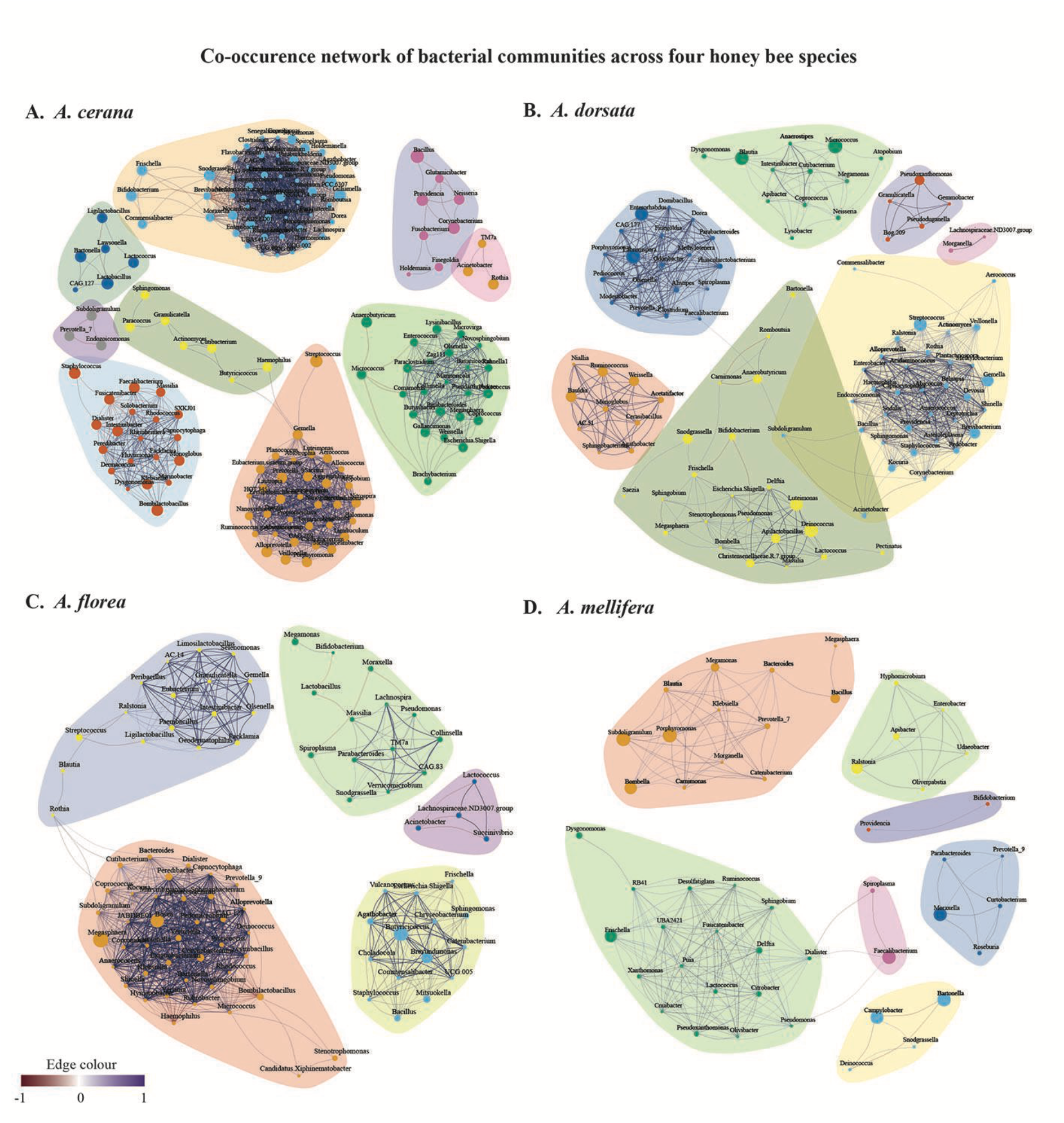
Structure of bacterial communities. Co-occurrence of bacterial microbiota across honey bee species: (A) *Apis cerana*, (B) *A. dorsata*, (C) *A. florea*, and (D) *A. mellifera*. In all these networks, the node size is the relative abundance of the bacterial genera, and the size and colour of the vertices denote the extent and direction of cooccurrence (i.e. positive or negative) between bacterial genera, respectively.

We further found that the fungal metacommunity network shows 11 modules, of which eight are isolated and without any intermodular connections, whereas only three modules are connected with only negative co-occurrence between four fungal genera *Excerohilum, Curvularia, Epicoccum, Malassezia* (Fig S10B). While comparing fungal co-occurrence networks across honeybee species, we found that *A. mellifera* hosts a fungal cobiont network which is highly modular and nested at the same time, whereas the connectance of the fungal network is highest in *A. florea* and *A. dorsata* harbours the least modular, connected and nested fungal network (Fig 5, Table S6). Similar to bacterial co-occurrence, fungal cobionts show species-specific community structures and associations. Moreover, we examined the co-occurrence meta-network between the most abundant bacterial and fungal cobionts from all four honeybee species. We found eight modules, out of which two modules are isolated and mostly composed of fungal genera (Fig S10C). In this microbial network, most of the intermodular connections are mediated by bacterial genera such as *Bombilactobacillus, Gilliamella, Carnimonas, Pseudomonas, Stenotrophomonas, Dysgonomonas, Commensalibacter, Atopobium, Cutibacterium, Granulicatella* and *Brevibacterium*. Among fungi, *Davidiella* and *Mucor* show the highest co-occurrence with bacterial communities (Fig S10C). Moreover, we compared co-occurrence patterns of microbes across honey bee species with continuity correction and found bacterial microbes in *A. florea* were significantly different compared to *A. dorsata* and *A. mellifera* (Table S7), whereas fungal co-occurrences differed only between *A. florea* and *A. dorsata* (Fig S8). Here, our results show that unlike fungi, bacterial co-occurrence networks formed highly species-specific communities, with core bacteria driving intermodular connectivity within the broader microbial metacommunity (i.e., general microbiome assemblages across the four honey bee species) (Fig. S10D).

**Figure 5.**
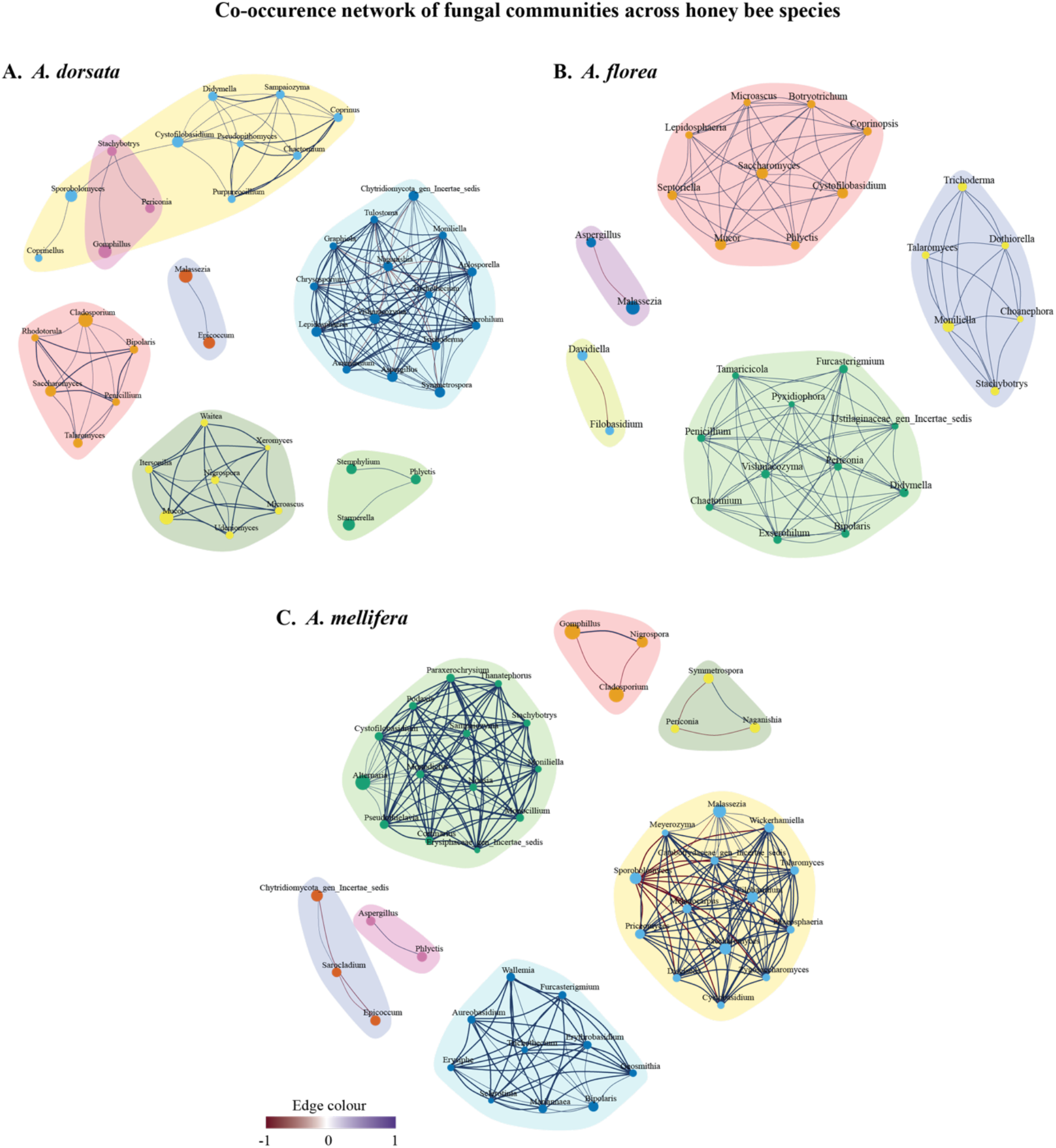
Structure of fungal communities. Co-occurrence of fungal microbiota across honey bee species: (A) *A. dorsata*, (B) *A. florea*, and (C) *A. mellifera*. In all these networks, the node size is the relative abundance of the fungal genera, and the size and colour of the vertices denote the extent and direction of cooccurrence (i.e. positive or negative) between fungal genera, respectively.

### Bayesian Inference indicates critical cobionts for microbiome stability and effects of perturbation on microbiome across honeybee species

To understand the stability of microbial networks in each species, we performed extinction rate simulations and found that bacterial communities in *A. cerana* have a net low rate of extinction because of the greater bacterial diversity and highly modular network. Conversely, bacterial networks in the other three honeybee species show a relatively higher extinction rate, which could be associated with less diverse and less modular bacterial networks hosted in them (Fig 6). In the case of the fungal network, we did not find differences in the extent of initial extinction across three honey bee species, and the rates of extinction were also not markedly different (Fig 6). Subsequently, to understand the directional co-occurrences within both bacterial and fungal metacommunities that may drive the varying stability, we implemented a Bayesian network approach and identified directional relationships among 34 bacterial (including *Apibacter, Porphyromonas, Bombella, Bartonella, Atopobium, Frischella, Rothia*) and fungal taxa (including *Nigrospora, Fusaria*) (Fig. S11), revealing the key microbial participants that may influence metacommunity and could shape the stability patterns observed in our extinction simulations. Our results show that while fungal networks exhibited uniform vulnerabilities across hosts, bacterial networks displayed host-specific stability, probably driven by a few keystone microbes.

**Figure 6.**
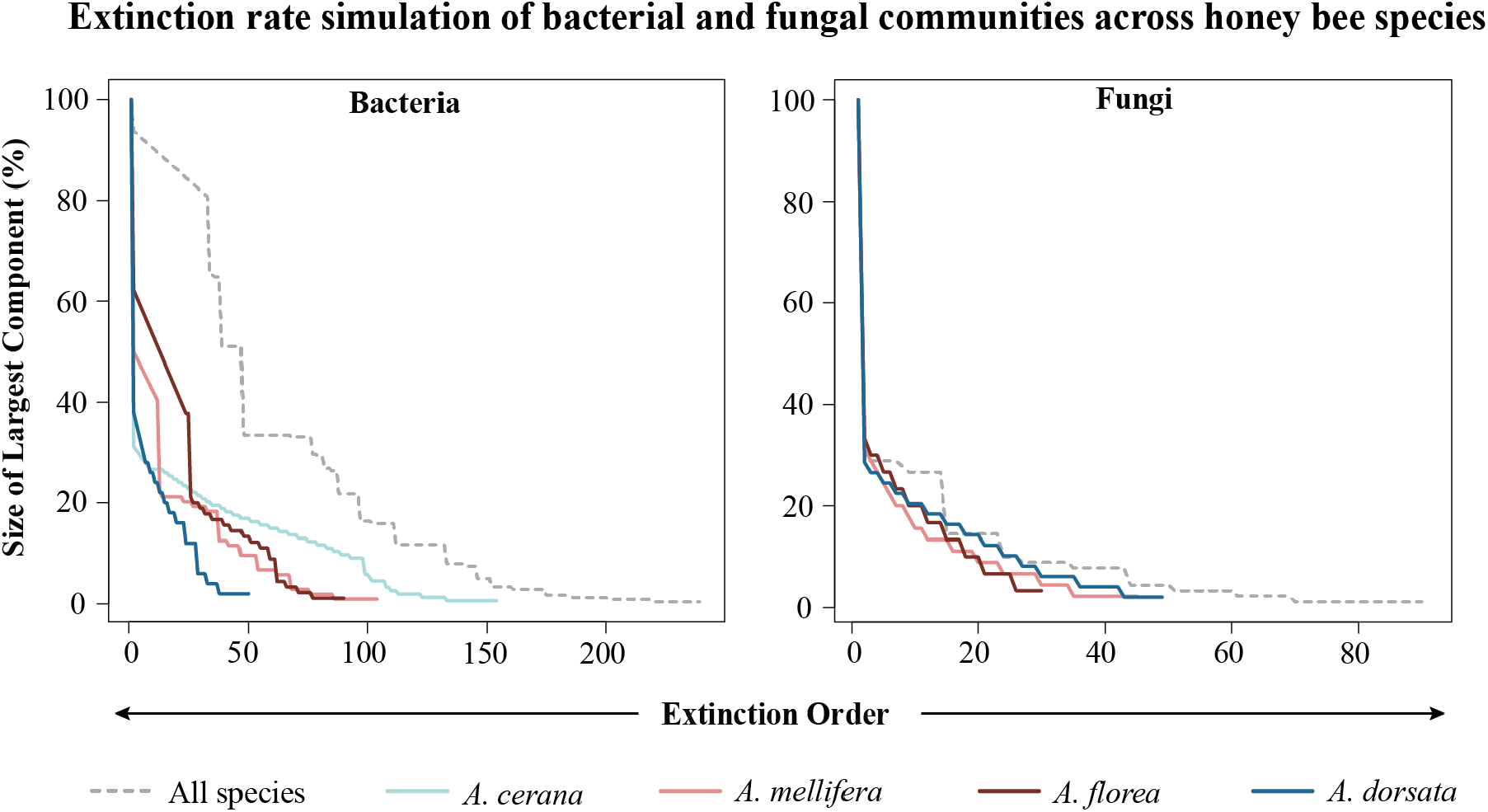
Extinction rate simulation. A comparative view of extinction rate simulation of bacterial (A) and fungal (B) communities to understand microbiome stability across honeybee species.

## DISCUSSION

In this study, we present a comparative framework that includes four major cooccurring *Apis* honeybee species, providing insights into how host physiology drives the assembly and structural stability of pollinator microbiomes, even when hosts share identical habitats. Using a simplified monoculture landscape, we controlled for the confounding effects of environmental heterogeneity in the pollination network and compared microbial sharing across four co-occurring *Apis* species. We show that, despite sharing a highly compartmentalised pollinator network (i.e., different honeybee species dominating at different field sites), microbial community assemblages were distinct across the four *Apis* species, characterised by strong host-mediated bacterial filtering. Our results also suggest that *the A. cerana* microbiome functionally diverged in lipid metabolism, antibiotic synthesis, and secondary metabolite metabolism, whereas the microbiomes of *A. florea* and *A. mellifera* have diverged metabolically in both carbohydrate and lipid metabolism (Fig S2, Dataset S4). Conversely, comparable fungal community compositions across species suggest no such divergence in fungal microbial composition. Using co-occurrence networks and extinction simulations, we demonstrate that these diverse bacterial community assemblies shape community stability. For example, the native *A. cerana* maintains a highly modular and diverse bacterial network that reduces microbial extinction rates, whereas the domesticated *A. mellifera* harbours a highly nested, less diverse bacterial network, making its microbial community more susceptible to structural collapse^25^. Overall, we predict that species-specific physiology, rather than resource homogenization, landscape use or varying foraging behaviour, shapes the structural stability of bacterial co-occurrence networks, which is in congruence with the previous studies on *A. mellifera* where it shows genetic background shapes gut microbiota^26^, highlighting the role of microbiome stability in pollinator communities^17,27,28^.

The spatial and behavioural partitioning observed in our foraging data illustrates how intrinsic host traits supersede environmental homogeneity to govern microbial acquisition. Although *A. mellifera* and *A. dorsata* functionally dominated the pollinator network in terms of visitation frequency and floral handling time, this did not correlate with higher microbial richness. Instead, the less dominant *A. cerana* harboured the highest bacterial alpha diversity. This uncoupling of foraging abundance from microbial richness contradicts the assumption that greater environmental exposure via floral contact strictly yields greater microbial diversity^29^. Rather, this incongruence indicates that microbial assembly could be regulated by species-specific physiological and behavioural traits, including colony size, intra-colonial grooming dynamics, and the stringency of vertical transmission pathways^11,30^. For instance, the reduced bacterial diversity observed in *A. mellifera* likely reflects the genetic bottlenecks and standardised rearing practices associated with domestication, which are known to constrain microbial acquisition and homogenise gut communities compared to wild populations^17,31^. Conversely, the higher microbial richness in native bees such as *A. cerana* may reflect greater physiological permissiveness or diverse social transmission routes that facilitate the retention of a broader array of environmentally acquired microbes. Our finding contradicts the previous finding of low microbial diversity in overwintering *A*. cerana when compared with *A. mellifera*, which could be due to different subspecies occurring in both regions showing divergence in their microbial community structure^32^. Studies expanding beyond the focal (social bee) taxa and examining the broader assemblage of insects visiting the same flowers, including solitary bees, butterflies, and other floral visitors, in order to assess the extent of microbial niche specialisation within a shared environmental context would be more insightful, explaining the ecological implications of microbial community interaction with pollinators.

Beyond absolute diversity, the compositional structure of these metacommunities elucidates profound differences in how these four species process environmental microbial inputs. Consistent with established models of *Apis* microbiomes, we identified a conserved core gut microbiota— including *Lactobacillus, Bifidobacterium, Snodgrassella*, and *Gilliamella*—that functions as a structural hubs interconnecting the broader metacommunity^18,19^. However, the unique microbial taxa acquired by each species expose varying degrees of environmental and anthropogenic effects. We detected human-associated gut taxa (e.g., *Faecalibacterium* in *A. mellifera*; *Fusobacterium* and *Neisseria* in *A. cerana*) and water-borne bacteria (e.g., *Endozoicomonas*), strongly indicating that agricultural water sources and proximity to human habitation serve as active conduits for horizontal microbial translocation into native pollinator networks. Furthermore, while bacterial taxa were strictly partitioned by host identity, fungal assemblages—including plant commensals such as *Aspergillus* and known agricultural pathogens such as *Daividiella* and *Alternaria*—were shared across all four bee species. This cross-species fungal homogenization supports the hypothesis that flowers serve as primary ecological hubs for horizontal fungal transmission^15,33^, whereas bacterial cobionts may undergo intense, host-mediated post-acquisition filtering. The differential presence of potentially pathogenic genera, such as *Spiroplasma* and *Paenibacillus* ^34,35^, further underscores that these native bees could experience distinct physiological and pathological outcomes within the same foraging landscape.

Interestingly, our study demonstrates that the topological architecture of these host-specific microbiomes directly dictates their susceptibility to ecological perturbation. The variation in structural modularity, connectance, and nestedness across the four species suggests differential vulnerabilities to both endogenous dysbiosis and exogenous environmental stressors^3^. Extinction rate simulations revealed that *A. cerana*, despite its susceptibility to initial microbial loss, maintains a significantly lower extinction rate at later stages due to its highly modular community structure, which prevents localised disturbances from cascading through the entire microbial communit^36^. Conversely, the highly nested and low-diversity microbial modules within *A. mellifera* show a higher systemic extinction rate at a later stage as well, indicating a fragile metacommunity that is highly susceptible to external perturbations, such as infection risk and insecticide-related dysbiosi^37^. Moreover, as *A. mellifera* is a non-native managed species in this landscape, microbiota composition may be further shaped by its management practices or by its inherent lower susceptibility to acquiring environmental microbiota, warranting further exploration. To identify the mechanistic drivers of this stability, our Bayesian network approach identified 34 specific microbial genera that act in a directional manner on stabilising the metacommunity^38^. These stabilising taxa could function as structural hubs, buffering the network against cascading effects of perturbation by maintaining critical metabolic redundancies and spatial modularity. Varying compositions of these keystone taxa in the core microbiomes of native versus managed bees provide a possible explanation for why the microbial community in *A. mellifera* exhibits a high extinction probability compared to the robust, compartmentalised microbiome composition of *A. cerana*. Consequently, identifying and protecting these microbial hubs will be critical for developing microbiome-directed interventions to mitigate pollinator decline.

In conclusion, the comparative metabarcoding approach utilised in this study provides a unique comparative framework for disentangling the complexities of host-microbiome biology in native paleotropical honey bees. While our methodology relies on amplicon-based sequencing—which inherently possesses lower functional resolution than whole-genome shotgun metagenomics and cannot definitively resolve spatial localisation within the host without tissue-specific sampling—it nonetheless robustly identifies the structural paradigms of microbial assembly. We have demonstrated that utilising a monoculture resource condition as an ecological background is instrumental in revealing how different honey bee species acquire, filter, and structure both beneficial and potentially adversarial microbes. Our integration of network topology, extinction simulations, and Bayesian modelling highlights that, rather than the effects of anthropogenic land use on the pollinator microbiome, species-specific network modularity acts as a primary defence against microbial collapse and could have greater implications for emerging infections in the pollinator network. Moving forward, integrating behavioural ecology with metacommunity network dynamics will be paramount to accurately assessing the ecological risks posed to globally declining pollinator populations.

## METHODS

### Study region and sampling sites

We conducted field sampling in mustard (*Brassica juncea*) fields located in the Rewari district of Haryana, northwestern India (28°09.302’, 76°22.459’). This semi-arid region, situated at an elevation of approximately 900 feet in the Aravalli foothills, comprises a heterogeneous matrix of forest patches, agricultural land, and human settlements. Mustard, an herbaceous annual in the Brassicaceae family, is the dominant winter cash crop in Haryana, sown between September and October, with flowering occurring from December to January. During this period, in northwestern India, mustard is cultivated over six lakh hectares, making a large seasonal monoculture agrolandscape. Beekeepers routinely position large numbers of *Apis mellifera* colonies at field margins, bringing managed populations into close contact with the three native species sampled in this study: *A. cerana, A. dorsata*, and *A. florea*. These species collectively represent a wide range of colony sizes, nesting substrates, and forager morphologies. In this monocultural system dominated by a single mass-flowering resource, shared floral visitation is expected to promote microbial exchange and homogenise microbial communities across species, thereby reducing the effects of preferential foraging of honey bee species on their microbiome^39^. Moreover, biological differences among hosts may impose species-specific filters on microbiome composition^40^. Seven fields were selected at least 1.5 km apart to ensure spatial independence. We further classified the bees based on their typical nesting habits (e.g., *A. florea* and *A. dorsata* are open nesters vs. *A. mellifera* and *A. cerana* are cavity nesters) along with vegetation cover, and surrounding land use across these field sites (details in *Exploring the land use variables* method section) provided a natural gradient to evaluate the relative contributions of host identity and shared foraging environment to microbiome structure (Fig. S1).

### Flower visitor observations and sampling bees

Each mustard field consisted of 100 meter transects covering both the central and peripheral regions of the field. Point counts (15-minute intervals) were conducted at 10-15-meter intervals along the transect (depending on the field size). During this 15-minute duration, we observed a patch of flowers and recorded the types and number of insect visitors. For a subset of these visitors (including the four species of honey bees), we also recorded how much time individuals from each species spent on each flower (Dataset S1). In total, visitor observations were carried out for 2820 minutes across seven sites. In each field, during the peak flowering season (December 2022-January 2023), bees foraging on the flowers were directly captured into sterile 15 ml tubes. These tubes were then added with 70% ethanol (AR) and transported to the laboratory, and temporarily stored at -80°C till DNA extraction.

### Extraction of DNA for microbial profiling

Six workers of each honeybee species (viz. *A. dorsata, A. cerana, A. florea*, and *A. mellifera*) from each of the seven survey fields were taken for DNA extraction^22^. During sample preparation, the heads of the bees were excluded to circumvent PCR inhibitors found in their compound eyes^41^. Additionally, wings and legs were removed, and the thorax and abdomen parts were washed with 70% ethanol to remove any contamination from the external body surfaces, such as pollen and plant remnants^41^. For each experimental sample for DNA extraction, we pooled these six individuals together (Dataset S2). We used the Wizard Genomic DNA Purification Kit from Promega and followed the manufacturer’s protocol for DNA extraction from microbial samples. The samples were then sequenced using the Illumina MiSeq platform 250 paired-end technique at miBiome Therapeutics LLP.

### Processing of amplicon sequencing data

The sequenced raw reads for bacteria and fungi were quality checked using FastQC, and primers were removed using cutadapt, and the paired reads were then quality filtered out (>30 phred scores). We have targeted V3–V4 and ITS1 region for characterising bacterial and fungal microbiome, respectively^42^. Unlike well-defined V3–V4 barcoding regions for bacterial reads, the ITS1 region used for fungal metabarcoding is flanked by conserved 18S and 5.8S regions. Therefore, to avoid sequence contamination, we used ITSxpress (v2.1.3) to extract the ITS1 region^43^. A standalone algorithm *DADA2* was implemented to infer Amplicon sequence variants (ASVs)^44^. The reads were further trimmed and truncated for quality check (including the range of read lengths: in case of bacteria, maximum 270 bp and minimum 220bp; in case of fungi, maximum 290 bp and minimum 210 bp) with the maximum of 2 bp expected errors^45^. An error model was generated for both the forward and reverse reads for denoising measures. Paired reads were merged, and a quality check was done for chimeric sequences. Finally, using the “assignTaxonomy” function from the DADA2 and databases such as SILVA, RDP, GTDB and Unite^46–49^ we assigned taxonomy for the bacterial reads and fungal reads. We further characterized bacterial ASVs to obtain functional genes using PICRUSt2 (Phylogenetic Investigation of Communities by Reconstruction of Unobserved States 2, v2.6.3)^24^. The predicted functional metagenomic profiles were further analysed using the “ggpicrust2” package (v1.7.4)^50^. Differential abundance analysis of KEGG pathways across bee species was performed using the ALDEx2 method, and p-values were adjusted for multiple testing using the Benjamini–Hochberg procedure^24^. Subsequently, we took the functional genes with adjusted p-value less than 0.05 and performed a canonical variate analysis to obtain differences between functional metaprofiles among the four honey bee species.

### Characterising the landscape heterogeneity and their effects on microbial profile

We have used the open-source geographic information system QGIS (v3.34) to characterise the land use types around the study sites. The UTM (Universal Transverse Mercator) region 43N with EPSG:32643 has been selected as a coordinate reference system for the geographic projection. Subsequently, the latitude and longitude of the fields were georeferenced on the Google satellite map^51^. We further used a polygon encompassing the coordinates of the fields to obtain the area of the field. The variables such as forest area, agricultural land, roads, human settlements, and secondary vegetation were marked to estimate the heterogeneity of the landscape. We extracted the proportion of all these variables within the 200-metre buffer using the intersection function from QGIS.

### Statistical analysis

a. <u>Analysis and visualization of relative abundance of honey bees and foraging behaviour</u>: We used Kruskal-Wallis tests to evaluate the differences in relative abundance and foraging time (measured as time of handling per flower) among the four bee species. Post hoc pairwise comparisons were conducted using Wilcoxon rank-sum tests. The network constructed between fields and bees (bipartite-plotweb).
b. <u>Estimation of relative abundance of microbiota and comparisons</u>: The reads classified as mitochondria and chloroplasts are excluded from the analysis, and the relative abundance of all the taxa and diversity indices was estimated using the R package “phyloseq”^52^. We used the Kruskal-Wallis test to evaluate differences in richness and alpha diversity of bacterial and fungal genera among bee species. Post hoc pairwise comparisons were conducted using Dunn’s tests. To analyse the overlap in bacterial and fungal genera composition among the four bee species at different sites, we used the Bray-Curtis dissimilarity distance measure and conducted a PERMANOVA (using the “adonis2” package in R) with 999 permutations and then visualized using principal coordinate analysis. To understand whether these microbial sharing patterns between honeybee species may drive more similarities in the microbial profile, we performed a homogeneity of variance test to estimate group dispersion.
c. <u>Estimation of co-occurrence networks of microbial communities and Bayesian inference on community structure</u>: Bacterial co-occurrence networks were estimated and visualised where network nodes and edges represented genera of ASVs^38^ and the co-occurrence properties between them, respectively, using the R-packages “corr.test” and “igraph”^53^ (for details see SI method). To understand directional relationships among microbial modules and to assess key taxa that influence the network structure, we used probabilistic graphical models to estimate directed acyclic graphs (DAGs). In these networks, links between microbial partners are directional, and their orientation is inferred from direct stochastic dependencies between pairs of nodes based on local probability distributions. Here, we chose a signed Bayesian network (sBN) approach^54^, a constraint-based algorithm better suited to causal inference than score-based approaches.
d. <u>Analysing the effects of biotic and abiotic parameters on microbial abundance</u>: We have used a generalised linear model (glm) to find the association between the relative abundance of each honeybee species with its microbial abundance. Similarly, we have used glm to identify any association of the foraging parameter (denoted by average time spent at flowers) of each honey bee species with its microbial abundance. Additionally, to investigate the effect of landscape use on bee species and microbial abundance, we have used a two-block partial least squares from the R package “Morpho” on relative microbial abundance and the habitat proportion table from QGIS (mentioned above) to estimate covariance.

## Acknowledgements

We also acknowledge the laboratory help from Pilot Dovih and Kripanjali Ghosh. We thank Rakesh Ahlawat and Manan Mehta for their support during field data collection.

## DATA AVAILABILITY

Data used in this manuscript are available in the supplementary material, and raw sequences are available upon request to the corresponding authors.

