## Supplementary Information for "TRACKING THE COMPOSITION AND STABILITY OF MICROBIOME ACROSS INDIAN SOCIAL HONEYBEES FORAGING IN A HOMOGENOUS RESOURCE LANDSCAPE"

**SUPPLEMENTARY METHODS**

**Estimation of co-occurrence networks of microbial communities and Bayesian inference on community structures:**

Bacterial co-occurrence networks were estimated and visualised where network nodes and edges represented genera of ASVs and the co-occurrence properties between them, respectively, using the R packages ”corr.test”^1^ and “igraph” ^2^. The following is a step-wise description of the network analysis.

i. The strength of the co-occurrence is computed as Pearson Correlation Coefficient and is reflected in the colouration of the edges (negative to positive correlation spectrum denoted as red to blue; Fig 4A-B) and their significance estimated using the "BH" adjustment on the p-values using *corr.test* function (represented by the width of the edges). All correlations with a P-adjusted value above 0.25 were not considered for the construction of the network, as well as in the signed Bayesian Network step (described below). The threshold of significance was set to a high value to account for influences which, while not significant (i.e. p<0.05), are still likely to have some effect on the interactions between ASV genera.
ii. Simultaneously, we used the mean of the normalised abundance of each genus of ASVs across all samples to represent the node size in the network. Next, we performed the Kruskal-Wallis test and used the -log(p-value) obtained from the test as the difference in the relative abundance of each ASV genus across honey bee species and represented as the node colour according to a heat scale (Fig S10A-B). The less significant the difference in abundance between two groups, the hotter the colour (yellower). The greater the significance of the difference, the cooler (bluer) the node’s colour.
iii. We used a force-directed *graphopt* algorithm to visualise the network ^2^, which relies on attractive and repulsive forces among nodes and subsequently simulates until it reaches an equilibrium. We used 500 iterations to reach the equilibrium.
iv. We have initially computed separate networks based on bacterial and fungal cooccurrence comprising all the honey bees. Subsequently, to assess the association between bacterial and fungal communities, we selected ASV genera that together account for 99% of the relative abundance of both bacteria and fungi and analysed their co-occurrence relationships. We further calculated bacterial and fungal co-occurrence networks for each species (where sample size was ≥3). We subsequently estimated graph-structure parameters, such as modularity, connectance, and nestedness, for each network to compare two co-occurrence networks across any two honey bee species.
v. To understand causal relations among microbial modules, we used probabilistic graphical models to estimate directed acyclic graphs (DAGs), in which each edge is directed and its direction is estimated from direct stochastic dependencies between nodes, represented as local probability distributions. Here, we chose a signed Bayesian network (sBN) approach, a constraint-based algorithm better suited to causal inference than score-based approaches. Note that sBN can only provide a partial DAG by calculating conditional independence and assuming the probability of independence as a proxy for graphical separation^3^.
vi. To optimise sBN inference for bacterial and fungal co-occurrences, we applied a threshold to select genera with sufficiently high abundance. We chose ASV genera, which constitute 99% of the relative abundance of both bacteria and fungi and analysed their cooccurrence relations.
vii. We used PC-stable, a constraint-based causality learning algorithm, to construct BNs, as it yields fewer false positives when estimating directionality, thereby increasing the sensitivity of Bayesian inference. We used the R package “bnlearn” to construct all the BNs^3^. The edges of the BNs were supported by the coefficient values from the co-occurrence networks calculated above, which transformed the BN into an sBN by distinguishing positive and negative correlations. A non-parametric bootstrap on 200 permuted inputs is computed to indicate the strength of directionality in each edge of the output network.

**SUPPLEMENTARY TABLES**

**Table S1**. Summary of the generalised linear model (GLM) analysis of honeybee abundance across species and fields

Model: Abundance ~ Species$\times$ Fields

| **Terms** | **Df** | | **Deviance** | **Residual df** | **Residual deviance** | **p value** |
| --- | --- | --- | --- | --- | --- | --- |
| Species | | 3 | 217.97 | 844 | 1170.8 | **< 0.001** |
| Field | | 6 | 86.03 | 838 | 1084.77 | **< 0.001** |
| Species ×Field | | 18 | 400.92 | 820 | 683.84 | **< 0.001** |

Pairwise contrast between honey bee species abundance across fields

| **Field** | **Contrast** | **Estimate** | **SE** | **z ratio** | **p value** |
| --- | --- | --- | --- | --- | --- |
| MUST1 | *A. cerana – A. dorsata* | -1.682 | 0.710 | -2.367 | 0.083 |
|  | *A. cerana – A. florea* | -1.099 | 0.715 | -1.537 | 0.415 |
|  | *A. cerana – A. mellifera* | -0.446 | 0.735 | -0.607 | 0.930 |
|  | *A. dorsata – A. florea* | 0.583 | 0.124 | 4.699 | **<0.001** |
|  | *A. dorsata – A. mellifera* | 1.236 | 0.211 | 5.847 | **<0.001** |
|  | *A. florea – A. mellifera* | 0.652 | 0.225 | 2.896 | **0.020** |
| MUST2 | *A. cerana – A. dorsata* | -0.833 | 0.213 | -3.903 | **0.001** |
|  | *A. cerana – A. florea* | -0.100 | 0.229 | -0.435 | 0.972 |
|  | *A. cerana – A. mellifera* | -1.368 | 0.210 | -6.525 | **<0.001** |
|  | *A. dorsata – A. florea* | 0.734 | 0.121 | 6.083 | **<0.001** |
|  | *A. dorsata – A. mellifera* | -0.535 | 0.079 | -6.794 | **<0.001** |
|  | *A. florea – A. mellifera* | -1.268 | 0.114 | -11.156 | **<0.001** |
| MUST3 | *A. cerana – A. dorsata* | -0.466 | 0.126 | -3.683 | **0.001** |
|  | *A. cerana – A. florea* | -0.698 | 0.124 | -5.649 | **<0.001** |
|  | *A. cerana – A. mellifera* | 0.453 | 0.204 | 2.224 | 0.117 |
|  | *A. dorsata – A. florea* | -0.233 | 0.104 | -2.232 | 0.115 |
|  | *A. dorsata – A. mellifera* | 0.918 | 0.192 | 4.773 | **<0.001** |
|  | *A. florea – A. mellifera* | 1.151 | 0.191 | 6.039 | **<0.001** |
| MUST4 | *A. cerana – A. dorsata* | 0.489 | 0.279 | 1.753 | 0.296 |
|  | *A. cerana – A. florea* | -1.009 | 0.169 | -5.984 | **<0.001** |
|  | *A. cerana – A. mellifera* | -0.928 | 0.166 | -5.590 | **<0.001** |
|  | *A. dorsata – A. florea* | -1.498 | 0.249 | -6.027 | **<0.001** |
|  | *A. dorsata – A. mellifera* | -1.417 | 0.247 | -5.741 | **<0.001** |
|  | *A. florea – A. mellifera* | 0.081 | 0.107 | 0.758 | 0.873 |
| MUST5 | *A. cerana – A. dorsata* | 0.930 | 0.267 | 3.483 | **0.003** |
|  | *A. cerana – A. florea* | 0.371 | 0.243 | 1.528 | 0.421 |
|  | *A. cerana – A. mellifera* | -0.590 | 0.115 | -5.136 | **<0.001** |
|  | *A. dorsata – A. florea* | -0.560 | 0.335 | -1.668 | 0.341 |
|  | *A. dorsata – A. mellifera* | -1.520 | 0.259 | -5.880 | **<0.001** |
|  | *A. florea – A. mellifera* | -0.961 | 0.233 | -4.121 | **0.002** |

| **Field** | **Contrast** | **Estimate** | **SE** | **z ratio** | **p value** |
| --- | --- | --- | --- | --- | --- |
| MUST6 | *A. cerana – A. dorsata* | 0.316 | 0.175 | 1.807 | 0.270 |
|  | *A. cerana – A. florea* | 0.557 | 0.209 | 2.668 | **0.038** |
|  | *A. cerana – A. mellifera* | -0.647 | 0.129 | -5.034 | **<0.001** |
|  | *A. dorsata – A. florea* | 0.242 | 0.222 | 1.086 | 0.698 |
|  | *A. dorsata – A. mellifera* | -0.963 | 0.149 | -6.438 | **<0.001** |
|  | *A. florea – A. mellifera* | -1.204 | 0.188 | -6.398 | **<0.001** |
| MUST7 | *A. cerana – A. dorsata* | -0.174 | 0.139 | -1.246 | 0.598 |
|  | *A. cerana – A. florea* | -0.029 | 0.187 | -0.155 | 0.999 |
|  | *A. cerana – A. mellifera* | -0.619 | 0.121 | -5.102 | **<0.001** |
|  | *A. dorsata – A. florea* | 0.145 | 0.186 | 0.777 | 0.865 |
|  | *A. dorsata – A. mellifera* | -0.446 | 0.121 | -3.696 | **0.001** |
|  | *A. florea – A. mellifera* | -0.590 | 0.173 | -3.412 | **0.004** |

Pairwise contrast between fields across honey bee species abundance

|  | **Species** | **Contrast** | **Estimate** | **SE** | **z ratio** | **p value** |
| --- | --- | --- | --- | --- | --- | --- |
|  | *A. cerana* | MUST1 - MUST2 | -0.780 | 0.736 | -1.060 | 0.940 |
|  |  | MUST1 – MUST3 | -0.974 | 0.714 | -1.364 | 0.821 |
|  |  | MUST1 – MUST4 | -0.671 | 0.723 | -0.929 | 0.968 |
|  |  | MUST1 – MUST5 | -1.064 | 0.713 | -1.491 | 0.750 |
|  |  | MUST1 – MUST6 | -1.027 | 0.716 | -1.435 | 0.783 |
|  |  | MUST1 – MUST7 | -0.887 | 0.714 | -1.243 | 0.877 |
|  |  | MUST2 – MUST3 | -0.194 | 0.228 | -0.851 | 0.979 |
|  |  | MUST2 – MUST4 | 0.109 | 0.253 | 0.431 | 1.000 |
|  |  | MUST2 – MUST5 | -0.284 | 0.225 | -1.262 | 0.869 |
|  |  | MUST2 – MUST6 | -0.247 | 0.232 | -1.063 | 0.939 |
|  |  | MUST2 – MUST7 | -0.107 | 0.227 | -0.472 | 0.999 |
|  |  | MUST3 – MUST4 | 0.303 | 0.180 | 1.682 | 0.628 |
|  |  | MUST3 – MUST5 | -0.090 | 0.138 | -0.650 | 0.995 |
|  |  | MUST3 – MUST6 | -0.053 | 0.150 | -0.354 | 1.000 |
|  |  | MUST3 – MUST7 | 0.087 | 0.141 | 0.613 | 0.996 |
|  |  | MUST4 – MUST5 | -0.393 | 0.176 | -2.228 | 0.281 |
|  |  | MUST4 – MUST5 | -0.356 | 0.186 | -1.915 | 0.470 |
|  |  | MUST4 – MUST7 | -0.216 | 0.179 | -1.208 | 0.892 |
|  |  | MUST5 - MUST6 | 0.037 | 0.146 | 0.252 | 1.000 |
|  |  | MUST5 – MUST7 | 0.177 | 0.137 | 1.292 | 0.856 |
|  |  | MUST6 – MUST7 | 0.140 | 0.149 | 0.940 | 0.966 |
|  |  | MUST1 - MUST2 | 0.068 | 0.093 | 0.739 | 0.990 |
|  |  | MUST1 – MUST3 | 0.242 | 0.102 | 2.370 | 0.211 |
|  |  | MUST1 – MUST4 | 1.499 | 0.245 | 6.111 | **<0.001** |
|  |  | MUST1 – MUST5 | 1.548 | 0.259 | 5.975 | **<0.001** |
|  | *A. dorsata* | MUST1 – MUST6 | 0.970 | 0.151 | 6.421 | **<0.001** |
|  |  | MUST1 – MUST7 | 0.621 | 0.119 | 5.198 | **<0.001** |
|  |  | MUST2 – MUST3 | 0.174 | 0.098 | 1.764 | 0.572 |
|  |  | MUST2 – MUST4 | 1.431 | 0.244 | 5.869 | **<0.001** |
| **Species** | | **Contrast** | **Estimate** | **SE** | **z ratio** | **p value** |
| *A. dorsata* | | MUST2 – MUST5 | 1.480 | 0.258 | 5.743 | **<0.001** |
|  |  | MUST2 – MUST6 | 0.902 | 0.149 | 6.068 | **<0.001** |
|  |  | MUST2 – MUST7 | 0.552 | 0.116 | 4.751 | **<0.001** |
|  |  | MUST3 – MUST4 | 1.257 | 0.248 | 5.077 | **<0.001** |
|  |  | MUST3 – MUST5 | 1.306 | 0.261 | 4.999 | **<0.001** |
|  |  | MUST3 – MUST6 | 0.728 | 0.155 | 4.704 | **0.001** |
|  |  | MUST3 – MUST7 | 0.379 | 0.124 | 3.053 | **0.037** |
|  |  | MUST4 – MUST5 | 0.049 | 0.344 | 0.142 | 1.000 |
|  |  | MUST4 – MUST6 | -0.529 | 0.272 | -1.949 | 0.448 |
|  |  | MUST4 – MUST7 | -0.879 | 0.255 | -3.441 | **0.010** |
|  |  | MUST5 - MUST6 | -0.578 | 0.284 | -2.035 | 0.393 |
|  |  | MUST5 – MUST7 | -0.927 | 0.269 | -3.453 | **0.010** |
|  |  | MUST6 – MUST7 | -0.349 | 0.167 | -2.096 | 0.355 |
| A. *florea* | | MUST1 - MUST2 | 0.219 | 0.146 | 1.497 | 0.747 |
|  |  | MUST1 – MUST3 | -0.574 | 0.126 | -4.560 | **0.001** |
|  |  | MUST1 – MUST4 | -0.582 | 0.130 | -4.465 | **0.002** |
|  |  | MUST1 – MUST5 | 0.405 | 0.246 | 1.645 | 0.653 |
|  |  | MUST1 – MUST6 | 0.629 | 0.205 | 3.067 | **0.035** |
|  |  | MUST1 – MUST7 | 0.182 | 0.189 | 0.964 | 0.962 |
|  |  | MUST2 – MUST3 | -0.793 | 0.125 | -6.322 | **<0.001** |
|  |  | MUST2 – MUST4 | -0.800 | 0.130 | -6.167 | **<0.001** |
|  |  | MUST2 – MUST5 | 0.187 | 0.246 | 0.758 | 0.989 |
|  |  | MUST2 – MUST6 | 0.410 | 0.205 | 2.002 | 0.413 |
|  |  | MUST2 – MUST7 | -0.037 | 0.189 | -0.194 | 1.000 |
|  |  | MUST3 – MUST4 | -0.008 | 0.106 | -0.075 | 1.000 |
|  |  | MUST3 – MUST5 | 0.979 | 0.235 | 4.172 | **0.001** |
|  |  | MUST3 – MUST6 | 1.202 | 0.191 | 6.308 | **<0.001** |
|  |  | MUST3 – MUST7 | 0.756 | 0.173 | 4.359 | **0.003** |
|  |  | MUST4 – MUST5 | 0.987 | 0.237 | 4.163 | **0.006** |
|  |  | MUST4 – MUST6 | 1.210 | 0.194 | 6.253 | **<0.001** |
|  |  | MUST4 – MUST7 | 0.764 | 0.177 | 4.324 | **0.003** |
|  |  | MUST5 - MUST6 | 0.223 | 0.285 | 0.783 | 0.987 |
|  |  | MUST5 – MUST7 | -0.223 | 0.274 | -0.815 | 0.984 |
|  |  | MUST6 – MUST7 | -0.446 | 0.237 | -1.882 | 0.493 |
| *A. mellifera* | | MUST1 - MUST2 | -1.702 | 0.206 | -8.276 | **<0.001** |
|  |  | MUST1 – MUST3 | -0.075 | 0.267 | -0.281 | 1.000 |
|  |  | MUST1 – MUST4 | -1.153 | 0.213 | -5.414 | **<0.001** |
|  |  | MUST1 – MUST5 | -1.208 | 0.211 | -5.734 | **<0.001** |
|  |  | MUST1 – MUST6 | -1.228 | 0.210 | -5.842 | **<0.001** |
|  |  | MUST1 – MUST7 | -1.060 | 0.212 | -5.002 | **<0.001** |
|  |  | MUST2 – MUST3 | 1.627 | 0.183 | 8.883 | **<0.001** |
|  |  | MUST2 – MUST4 | 0.549 | 0.087 | 6.297 | **<0.001** |
|  |  | MUST2 – MUST5 | 0.494 | 0.082 | 6.067 | **<0.001** |
|  |  | MUST2 – MUST6 | 0.474 | 0.080 | 5.901 | **<0.001** |

| **Species** | **Contrast** | **Estimate** | **SE** | **z ratio** | **p value** |
| --- | --- | --- | --- | --- | --- |
| *A. mellifera* | MUST2 – MUST7 | 0.642 | 0.085 | 7.553 | **<0.001** |
|  | MUST3 – MUST4 | -1.078 | 0.191 | -5.635 | **<0.001** |
|  | MUST3 – MUST5 | -1.133 | 0.189 | -6.003 | **<0.001** |
|  | MUST3 – MUST6 | -1.153 | 0.188 | -6.125 | **<0.001** |
|  | MUST3 – MUST7 | -0.985 | 0.190 | -5.180 | **<0.001** |
|  | MUST4 – MUST5 | -0.055 | 0.098 | -0.560 | 0.998 |
|  | MUST4 – MUST6 | -0.075 | 0.097 | -0.771 | 0.988 |
|  | MUST4 – MUST7 | 0.092 | 0.101 | 0.912 | 0.971 |
|  | MUST5 - MUST6 | -0.020 | 0.092 | -0.218 | 1.000 |
|  | MUST5 – MUST7 | 0.147 | 0.096 | 1.530 | 0.727 |
|  | MUST6 – MUST7 | 0.167 | 0.095 | 1.756 | 0.578 |

**Table S2**. Summary of the generalised linear model (GLM) for time spent by honey bees across species and fields

Model: Time spent ~ Species$\times$ Fields

| **Terms** | **Df** | | **Deviance** | **Residual df** | **Residual deviance** | **F statistic** | **p value** |
| --- | --- | --- | --- | --- | --- | --- | --- |
| Species | | 3 | 72013 | 2955 | 1170.8 | 106.163 | **< 0.001** |
| Fields | | 6 | 41452 | 2949 | 1084.77 | 30.555 | **< 0.001** |
| Species × Fields | | 18 | 30807 | 2931 | 683.84 | 7.569 | **< 0.001** |

Pairwise contrast between honey bee species across fields for time spent

| **Field** | **Contrast** | **Estimate** | **SE** | **t ratio** | **p value** |
| --- | --- | --- | --- | --- | --- |
| MUST1 | *A. cerana – A. dorsata* | -14.663 | 10.70 | -1.371 | 0.518 |
|  | *A. cerana – A. florea* | 3.679 | 10.80 | 0.342 | 0.986 |
|  | *A. cerana – A. mellifera* | 9.458 | 11.10 | 0.855 | 0.828 |
|  | *A. dorsata – A. florea* | 18.342 | 1.99 | 9.215 | **<0.001** |
|  | *A. dorsata – A. mellifera* | 24.121 | 3.27 | 7.377 | **<0.001** |
|  | *A. florea – A. mellifera* | 5.780 | 3.480 | 1.661 | 0.345 |
| MUST2 | *A. cerana – A. dorsata* | -0.013 | 3.350 | -1.371 | 1.000 |
|  | *A. cerana – A. florea* | -8.845 | 3.630 | 0.342 | 0.071 |
|  | *A. cerana – A. mellifera* | 7.645 | 3.300 | 0.855 | 0.095 |
|  | *A. dorsata – A. florea* | -8.832 | 1.970 | 9.215 | **<0.001** |
|  | *A. dorsata – A. mellifera* | 7.658 | 1.260 | 7.377 | **<0.001** |
|  | *A. florea – A. mellifera* | 16.490 | 1.870 | 1.661 | **<0.001** |
| MUST3 | *A. cerana – A. dorsata* | 0.778 | 1.970 | 0.395 | 0.979 |
|  | *A. cerana – A. florea* | 0.669 | 1.990 | 0.336 | 0.987 |
|  | *A. cerana – A. mellifera* | 8.391 | 3.050 | 2.750 | **0.031** |
|  | *A. dorsata – A. florea* | -0.109 | 1.710 | -0.064 | 1.000 |
|  | *A. dorsata – A. mellifera* | 7.614 | 2.880 | 2.647 | **0.041** |
|  | *A. florea – A. mellifera* | 7.722 | 2.890 | 2.671 | **0.038** |
| MUST4 | *A. cerana – A. dorsata* | -6.940 | 4.310 | -1.611 | 0.373 |
|  | *A. cerana – A. florea* | 2.956 | 2.620 | 1.126 | 0.673 |
|  | *A. cerana – A. mellifera* | 7.087 | 2.560 | 2.764 | 0.029 |
|  | *A. dorsata – A. florea* | 9.896 | 3.860 | 2.561 | 0.051 |
|  | *A. dorsata – A. mellifera* | 14.027 | 3.820 | 3.669 | **0.001** |
|  | *A. florea – A. mellifera* | 4.132 | 1.720 | 2.409 | 0.076 |
| MUST5 | *A. cerana – A. dorsata* | -13.511 | 3.960 | -3.415 | **0.004** |
|  | *A. cerana – A. florea* | -0.299 | 3.860 | -0.077 | 1.000 |
|  | *A. cerana – A. mellifera* | 2.592 | 1.870 | 1.385 | 0.509 |
|  | *A. dorsata – A. florea* | 13.212 | 5.090 | 2.598 | **0.047** |
|  | *A. dorsata – A. mellifera* | 16.103 | 3.800 | 4.236 | **0.001** |
|  | *A. florea – A. mellifera* | 2.890 | 3.700 | 0.781 | 0.863 |
| MUST6 | *A. cerana – A. dorsata* | -4.095 | 2.630 | -1.559 | 0.403 |
|  | *A. cerana – A. florea* | -6.377 | 3.140 | -2.031 | 0.177 |
|  | *A. cerana – A. mellifera* | 2.917 | 1.950 | 1.495 | 0.440 |
|  | *A. dorsata – A. florea* | -2.282 | 3.340 | -0.683 | 0.904 |
|  | *A. dorsata – A. mellifera* | 7.012 | 2.260 | 3.097 | **0.011** |
|  | *A. florea – A. mellifera* | 9.294 | 2.840 | 3.270 | **0.006** |
| MUST7 | *A. cerana – A. dorsata* | -0.569 | 2.100 | -0.271 | 0.993 |
|  | *A. cerana – A. florea* | -4.665 | 2.840 | -1.643 | 0.354 |
|  | *A. cerana – A. mellifera* | 7.049 | 1.840 | 3.834 | **0.001** |
|  | *A. dorsata – A. florea* | -4.097 | 2.820 | -1.455 | 0.465 |
|  | *A. dorsata – A. mellifera* | 7.618 | 1.800 | 4.224 | **0.001** |
|  | *A. florea – A. mellifera* | 11.714 | 2.630 | 4.454 | **0.001** |

Pairwise contrast between fields for time spent across species

| **Species** | **Contrast** | **Estimate** | **SE** | **t ratio** | **p value** |
| --- | --- | --- | --- | --- | --- |
| *A. cerana* | MUST1 - MUST2 | 5.909 | 11.100 | 0.532 | 0.998 |
|  | MUST1 – MUST3 | 6.685 | 10.700 | 0.622 | 0.996 |
|  | MUST1 – MUST4 | 4.616 | 10.900 | 0.424 | 1.000 |
|  | MUST1 – MUST5 | 9.188 | 10.700 | 0.855 | 0.979 |
|  | MUST1 – MUST6 | 10.377 | 10.800 | 0.964 | 0.962 |
|  | MUST1 – MUST7 | 7.550 | 10.700 | 0.703 | 0.992 |
|  | MUST2 – MUST3 | 0.776 | 3.570 | 0.217 | 1.000 |
|  | MUST2 – MUST4 | -1.293 | 3.940 | -0.328 | 1.000 |
|  | MUST2 – MUST5 | 3.278 | 3.550 | 0.922 | 0.969 |
|  | MUST2 – MUST6 | 4.468 | 3.620 | 1.236 | 0.880 |
|  | MUST2 – MUST7 | 1.641 | 3.540 | 0.463 | 0.999 |
|  | MUST3 – MUST4 | -2.069 | 2.780 | -0.745 | 0.990 |
|  | MUST3 – MUST5 | 2.503 | 2.190 | 1.141 | 0.916 |
|  | MUST3 – MUST6 | 3.692 | 2.290 | 1.611 | 0.675 |
|  | MUST3 – MUST7 | 0.865 | 2.170 | 0.398 | 1.000 |
|  | MUST4 – MUST5 | 4.571 | 2.760 | 1.657 | 0.645 |
|  | MUST4 – MUST6 | 5.760 | 2.840 | 2.030 | 0.396 |
|  | MUST4 – MUST7 | 2.934 | 2.740 | 1.070 | 0.937 |
|  | MUST5 - MUST6 | 1.189 | 2.270 | 0.524 | 0.999 |
|  | MUST5 – MUST7 | -1.638 | 2.150 | -0.762 | 0.988 |
|  | MUST6 – MUST7 | -2.827 | 2.250 | -1.257 | 0.871 |
| *A. dorsata* | MUST1 - MUST2 | 20.560 | 1.500 | 13.722 | **<0.001** |
|  | MUST1 – MUST3 | 22.125 | 1.640 | 13.484 | **<0.001** |
|  | MUST1 – MUST4 | 12.339 | 3.820 | 3.233 | **0.021** |
|  | MUST1 – MUST5 | 10.339 | 3.820 | 2.709 | 0.096 |
|  | MUST1 – MUST6 | 20.945 | 2.320 | 9.029 | **<0.001** |
|  | MUST1 – MUST7 | 21.644 | 1.840 | 11.732 | **<0.001** |
|  | MUST2 – MUST3 | 1.566 | 1.550 | 1.011 | 0.952 |
|  | MUST2 – MUST4 | -8.220 | 3.780 | -2.176 | 0.309 |
|  | MUST2 – MUST5 | -10.220 | 3.780 | -2.705 | 0.097 |
|  | MUST2 – MUST6 | 0.385 | 2.260 | 0.171 | 1.000 |
|  | MUST2 – MUST7 | 1.085 | 1.760 | 0.615 | 0.996 |
|  | MUST3 – MUST4 | -9.786 | 3.840 | -2.550 | 0.142 |
|  | MUST3 – MUST5 | -11.786 | 3.840 | -3.072 | **0.035** |
|  | MUST3 – MUST6 | -1.180 | 2.350 | -0.502 | 0.999 |
|  | MUST3 – MUST7 | -0.481 | 1.890 | -0.255 | 1.000 |
|  | MUST4 – MUST5 | -2.000 | 5.160 | -0.388 | 1.000 |
|  | MUST4 – MUST6 | 8.605 | 4.170 | 2.062 | 0.376 |
|  | MUST4 – MUST7 | 9.305 | 3.930 | 2.368 | 0.212 |
|  | MUST5 - MUST6 | 10.605 | 4.170 | 2.542 | 0.145 |
|  | MUST5 – MUST7 | 11.305 | 3.930 | 2.878 | 0.061 |
|  | MUST6 – MUST7 | 0.699 | 2.500 | 0.280 | 1.000 |

| *A. florea* | MUST1 - MUST2 | -6.615 | 2.360 | -2.797 | 0.076 |
| --- | --- | --- | --- | --- | --- |
|  | MUST1 – MUST3 | 3.675 | 2.050 | 1.793 | 0.553 |
|  | MUST1 – MUST4 | 3.893 | 2.080 | 1.874 | 0.498 |
|  | MUST1 – MUST5 | 5.210 | 3.910 | 1.334 | 0.836 |
|  | MUST1 – MUST6 | 0.321 | 3.120 | 0.103 | 1.000 |
|  | MUST1 – MUST7 | -0.794 | 2.910 | -0.272 | 1.000 |
|  | MUST2 – MUST3 | 10.289 | 2.100 | 4.902 | **<0.001** |
|  | MUST2 – MUST4 | 10.508 | 2.130 | 4.939 | **<0.001** |
|  | MUST2 – MUST5 | 11.825 | 3.930 | 3.007 | **0.042** |
|  | MUST2 – MUST6 | 6.936 | 3.160 | 2.197 | 0.297 |
|  | MUST2 – MUST7 | 5.821 | 2.950 | 1.974 | 0.432 |
|  | MUST3 – MUST4 | 0.219 | 1.770 | 0.123 | 1.000 |
|  | MUST3 – MUST5 | 1.536 | 3.750 | 0.409 | 1.000 |
|  | MUST3 – MUST6 | -3.353 | 2.930 | -1.145 | 0.914 |
|  | MUST3 – MUST7 | -4.469 | 2.700 | -1.653 | 0.647 |
|  | MUST4 – MUST5 | 1.317 | 3.770 | 0.350 | 1.000 |
|  | MUST4 – MUST6 | -3.572 | 2.950 | -1.212 | 0.890 |
|  | MUST4 – MUST7 | -4.687 | 2.720 | -1.720 | 0.602 |
|  | MUST5 - MUST6 | -4.889 | 4.430 | -1.104 | 0.927 |
|  | MUST5 – MUST7 | -6.004 | 4.280 | -1.401 | 0.801 |
|  | MUST6 – MUST7 | -1.115 | 3.590 | -0.311 | 1.000 |
| *A. mellifera* | MUST1 - MUST2 | 4.096 | 3.170 | 1.293 | 0.855 |
|  | MUST1 – MUST3 | 5.617 | 4.030 | 1.393 | 0.806 |
|  | MUST1 – MUST4 | 2.245 | 3.280 | 0.685 | 0.993 |
|  | MUST1 – MUST5 | 2.321 | 3.250 | 0.714 | 0.992 |
|  | MUST1 – MUST6 | 3.835 | 3.230 | 1.187 | 0.899 |
|  | MUST1 – MUST7 | 5.141 | 3.250 | 1.583 | 0.693 |
|  | MUST2 – MUST3 | 1.521 | 2.730 | 0.557 | 0.998 |
|  | MUST2 – MUST4 | -1.851 | 1.390 | -1.333 | 0.837 |
|  | MUST2 – MUST5 | -1.775 | 1.330 | -1.337 | 0.834 |
|  | MUST2 – MUST6 | -0.261 | 1.280 | -0.204 | 1.000 |
|  | MUST2 – MUST7 | 1.045 | 1.320 | 0.793 | 0.986 |
|  | MUST3 – MUST4 | -3.372 | 2.860 | -1.180 | 0.902 |
|  | MUST3 – MUST5 | -3.297 | 2.830 | -1.166 | 0.907 |
|  | MUST3 – MUST6 | -1.782 | 2.800 | -0.635 | 0.996 |
|  | MUST3 – MUST7 | -0.477 | 2.820 | -0.169 | 1.000 |
|  | MUST4 – MUST5 | 0.076 | 1.570 | 0.048 | 1.000 |
|  | MUST4 – MUST6 | 1.590 | 1.530 | 1.042 | 0.944 |
|  | MUST4 – MUST7 | 2.896 | 1.560 | 1.856 | 0.510 |
|  | MUST5 - MUST6 | 1.515 | 1.470 | 1.030 | 0.947 |
|  | MUST5 – MUST7 | 2.820 | 1.510 | 1.873 | 0.499 |
|  | MUST6 – MUST7 | 1.305 | 1.460 | 0.894 | 0.974 |

**Table** **S3**. Summary of the generalised linear mixed model (GLMM) analysis of bacterial genera richness across species

Model: Abundance ~ Species + (1|Fields); Family: Poisson (log link)

| **Community** | **Chisq** | **Df** | **P-value** |
| --- | --- | --- | --- |
| Bacteria | 104.137 | 3 | **<0.001** |
| Fungi | 43.952 | 3 | **<0.001** |

Post hoc comparisons of bacterial and fungal genera richness across honey bee species

| **Community** | **contrast** | **estimate** | **SE** | **z.ratio** | **p.value** |
| --- | --- | --- | --- | --- | --- |
| Bacteria | *A. cerana - A. dorsata* | 0.132 | 0.692 | 4.345 | **<0.001** |
|  | *A. cerana - A. florea* | 0.629 | 0.852 | 4.887 | **<0.001** |
|  | *A. cerana - A. mellifera* | 0.932 | 1.338 | 7.608 | **<0.001** |
|  | *A. dorsata - A. florea* | 0.496 | 0.160 | 0.996 | 0.751 |
|  | *A. dorsata - A. mellifera* | 0.800 | 0.646 | 3.983 | **<0.001** |
|  | *A. florea - A. mellifera* | 0.304 | 0.486 | 2.745 | 0.031 |
| Fungi | *A. cerana - A. dorsata* | -1.514 | -0.128 | -0.543 | 0.948 |
|  | *A. cerana - A. florea* | -1.157 | -0.059 | -0.241 | 0.995 |
|  | *A. cerana - A. mellifera* | -1.386 | -0.288 | -1.207 | 0.622 |
|  | *A. dorsata - A. florea* | 0.357 | 0.069 | 0.443 | 0.971 |
|  | *A. dorsata - A. mellifera* | 0.128 | -0.160 | -1.094 | 0.693 |
|  | *A. florea - A. mellifera* | -0.229 | -0.229 | -1.429 | 0.481 |

**Table S4.** Canonical variate statistics of pairwise comparisons between functional metaprofiles of bacterial cobionts across four honey bee species

| **Comparison** | **Mahalanobis**  **Distance** | **Probability** |
| --- | --- | --- |
| *A. cerana — A. dorsata* | 19.427 | **0.001** |
| *A. cerana — A. florea* | 25.012 | **<0.001** |
| *A. cerana — A. mellifera* | 22.766 | **<0.001** |
| *A. dorsata — A. florea* | 12.046 | **0.028** |
| *A. dorsata — A. mellifera* | 6.525 | 0.289 |
| *A. florea — A. mellifera* | 12.063 | **0.031** |

**Table** **S5**. Proportion of all the heterogeneous variable neighbouring the study sites

| **Sr no** | **Fields** | **Field area** | **Agricultural lands** | **Forest area** | **Human settlements** | **Roads** | **Secondary vegetation** |
| --- | --- | --- | --- | --- | --- | --- | --- |
| 1 | MUST1 | 0.05 | 0.45 | 0.27 | 0.19 | 0.04 | — |
| 2 | MUST2 | 0.08 | 0.46 | 0.45 | — | 0.02 | — |
| 3 | MUST3 | 0.09 | 0.78 | 0.11 | 0.01 | 0.01 | — |
| 4 | MUST4 | 0.06 | 0.89 | — | 0.02 | 0.04 | — |
| 5 | MUST5 | 0.04 | 0.82 | — | 0.01 | 0.03 | 0.09 |
| 6 | MUST6 | 0.06 | 0.92 | — | 0.02 | 0.01 | — |
| 7 | MUST7 | 0.05 | 0.93 | — | — | 0.02 | — |

**Table S6.** Network properties for bacterial and fungal community network across honey bee species

| **Microbial components** | **Network Properties** | **All species** | ***A. cerana*** | ***A. dorsata*** | ***A. florea*** | ***A. mellifera*** |
| --- | --- | --- | --- | --- | --- | --- |
| **Bacteria** | **Modularity** | 0.55 | 0.64 | 0.63 | 0.43 | 0.56 |
|  | **Connectance** | 0.15 | 0.17 | 0.15 | 0.23 | 0.18 |
|  | **Nestedness** | 22.79 | 22.00 | 25.33 | 31.98 | 32.03 |
| **Fungi** | **Modularity** | 0.87 | — | 0.57 | 0.61 | 0.67 |
|  | **Connectance** | 0.08 | — | 0.17 | 0.25 | 0.20 |
|  | **Nestedness** | 29.26 | — | 33.76 | 39.11 | 41.97 |
| **Bacteria and fungi** | **Modularity** | 0.73 | — | — | — | — |
|  | **Connectance** | 0.13 | — | — | — | — |
|  | **Nestedness** | 29.24 | — | — | — | — |

**Table** **S7.** Comparisons of bacterial genera across honey bee species using the pairwise Wilcoxon rank sum test with continuity correction.

|  | *A. cerana* | *A. dorsata* | *A. florea* |
| --- | --- | --- | --- |
| *A. dorsata* | 1.00 | — | — |
| *A. florea* | 1.00 | 0.01 | — |
| *A. mellifera* | 1.00 | 1.00 | 0.13 |

**Table** **S8.** Comparisons of fungal genera across honey bee species using the pairwise Wilcoxon rank sum test with continuity correction.

|  | *A. cerana* | *A. dorsata* | *A. florea* |
| --- | --- | --- | --- |
| *A. dorsata* | 1.00 | — | — |
| *A. florea* | 1.00 | 0.58 | — |
| *A. mellifera* | 1.00 | 0.03 | 1.00 |

**SUPPLEMENTARY FIGURES**

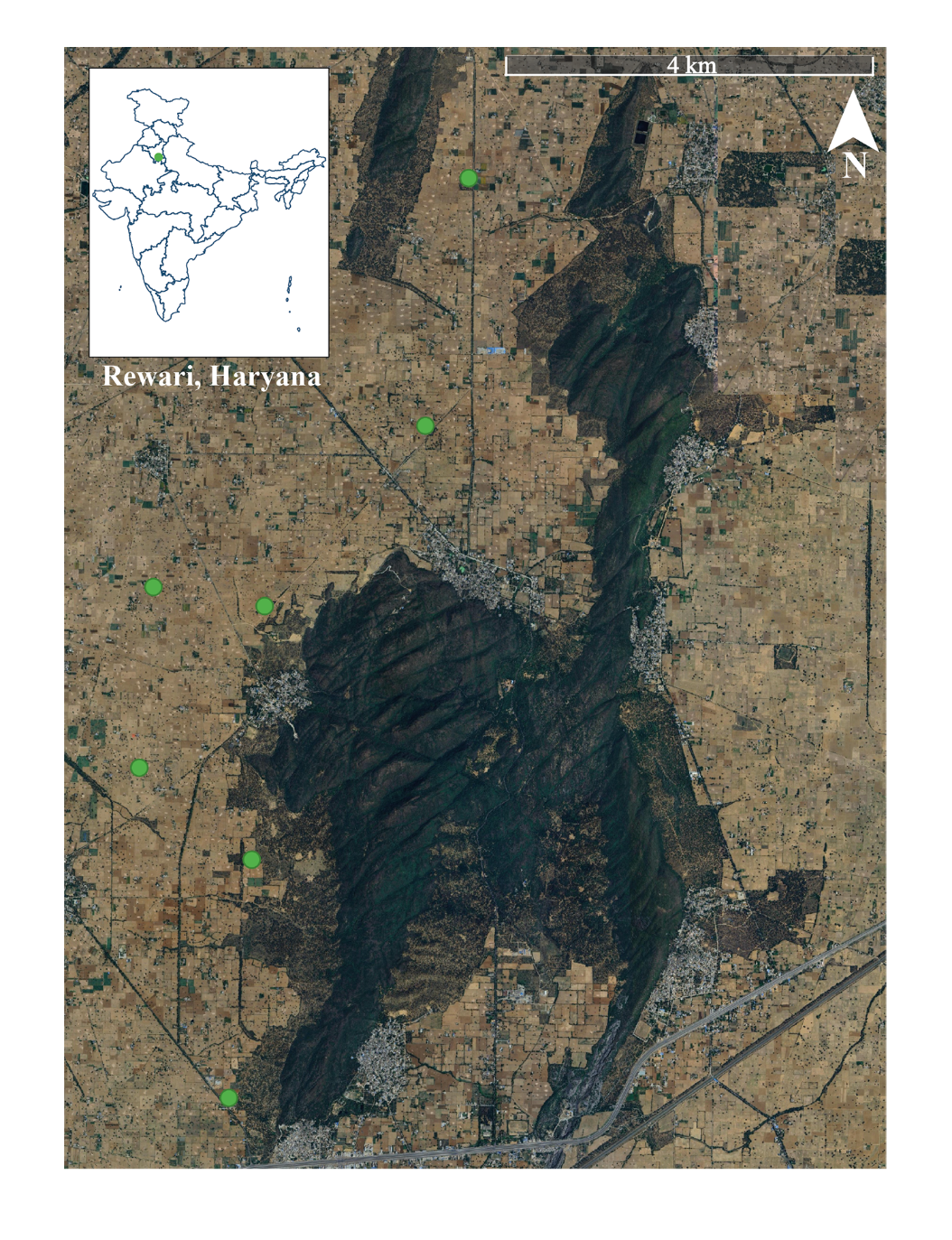

**Fig S1**. Map of the study area located in the semi-arid foothills of the Aravalli Range (Rewari, Haryana). Green dots indicate seven sampling sites (MUST1 – MUST7).

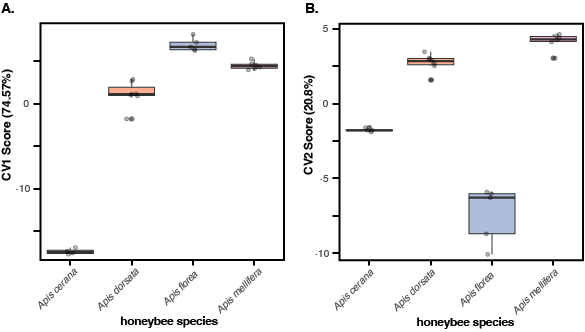

**Fig S2. Comparison of the functional metaprofile of the honeybee microbiome based on canonical variate analysis on the estimated abundance of significantly divergent functional KEGG pathways.** Both (A) canonical variate axis 1 (CV1) and (B) canonical variate axis 2 (CV2) show divergences in the functional profile of microbes across honey bee species in two orthogonal dimensions of variance.

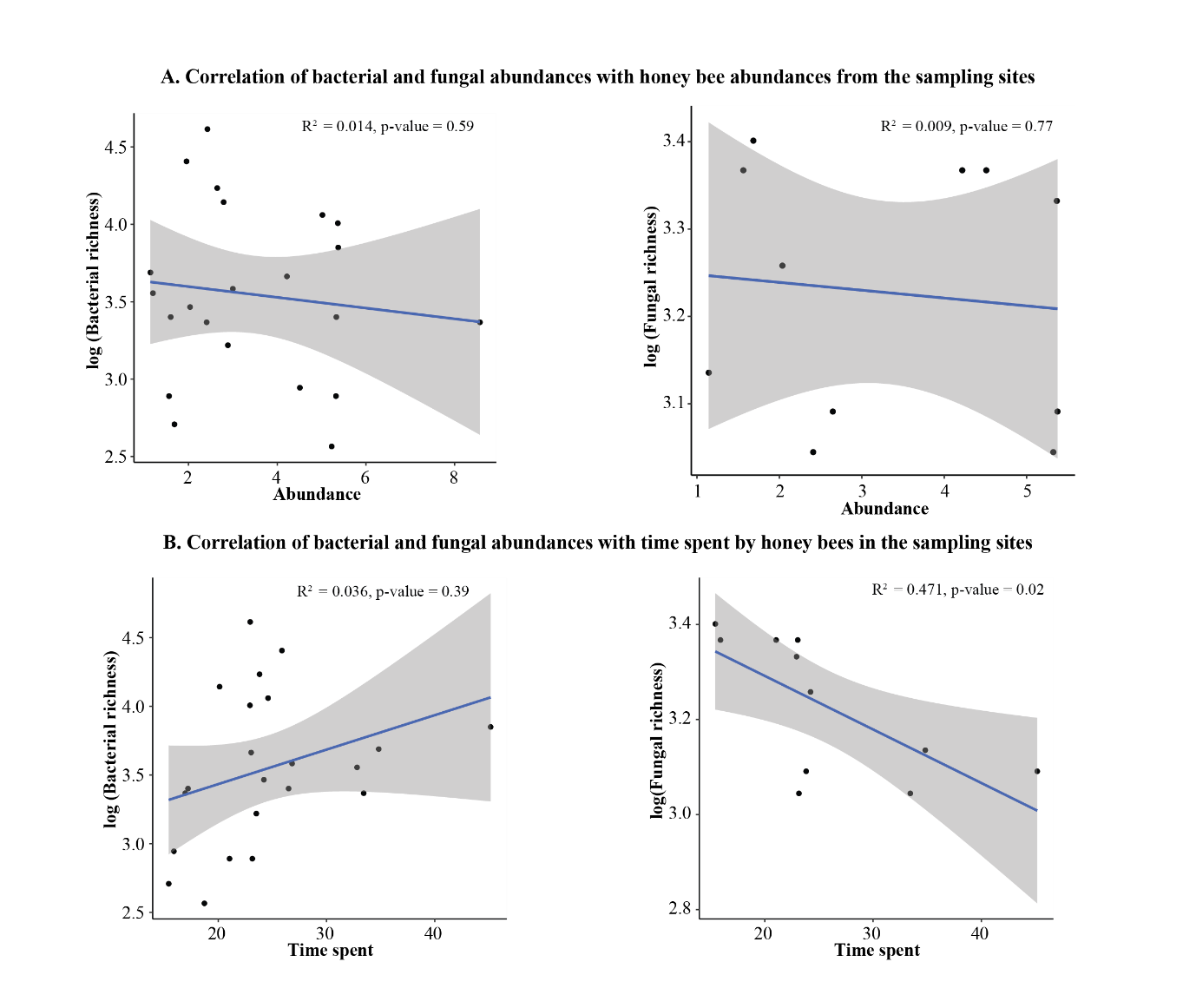

**Fig S3**. **Correlation of bacterial and fungal richness with honey bee abundance and time spent on the sampling sites.** (A) Relationship between honey bee abundance and microbial richness across sampling sites. The left panel shows the correlation between abundance and log-transformed bacterial richness, while the right panel shows the correlation between abundance and log-transformed fungal richness. (B) Relationship between time spent by honey bees on sites and microbial richness. The left panel represents the correlation between time spent and log-transformed bacterial richness, and the right panel represents the correlation between time spent and log-transformed fungal richness. In all panels, dots represent individual samples from the sampling sites, the solid line indicates the fitted regression trend, and the shaded region represents 95% confidence interval around the regression line. Reported R^2^ values correspond to multiple R-squared value from the linear regression, along with associated p-values.

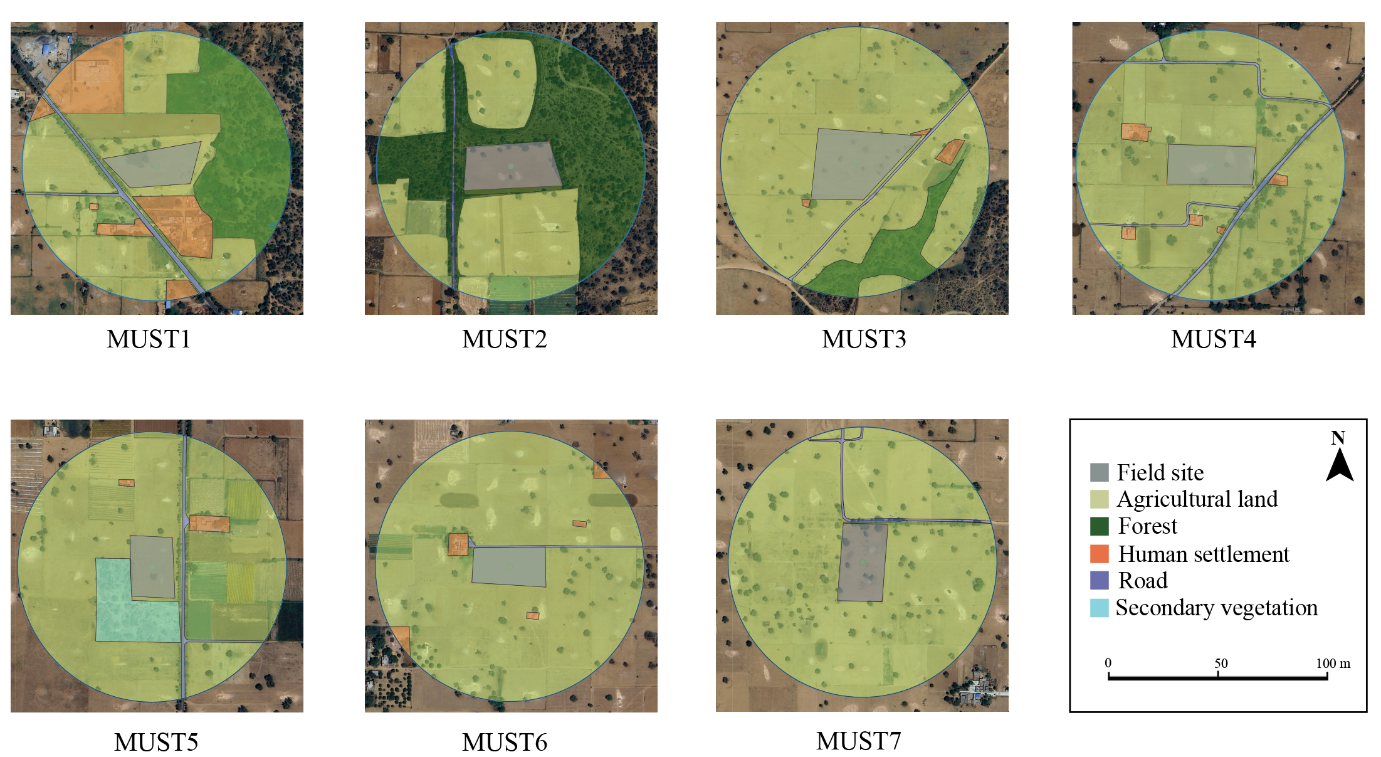

**Fig S4. Spatial representation of land use composition surrounding field sites (MUST 1-7).** In each field site, a 200 m radius buffer is represented by blue circle within which polygons indicate different land-use types (agricultural land (light yellow), forest (dark green), human settlements (orange), roads (blue), and secondary vegetation (sky blue)) quantified for subsequent analysis.

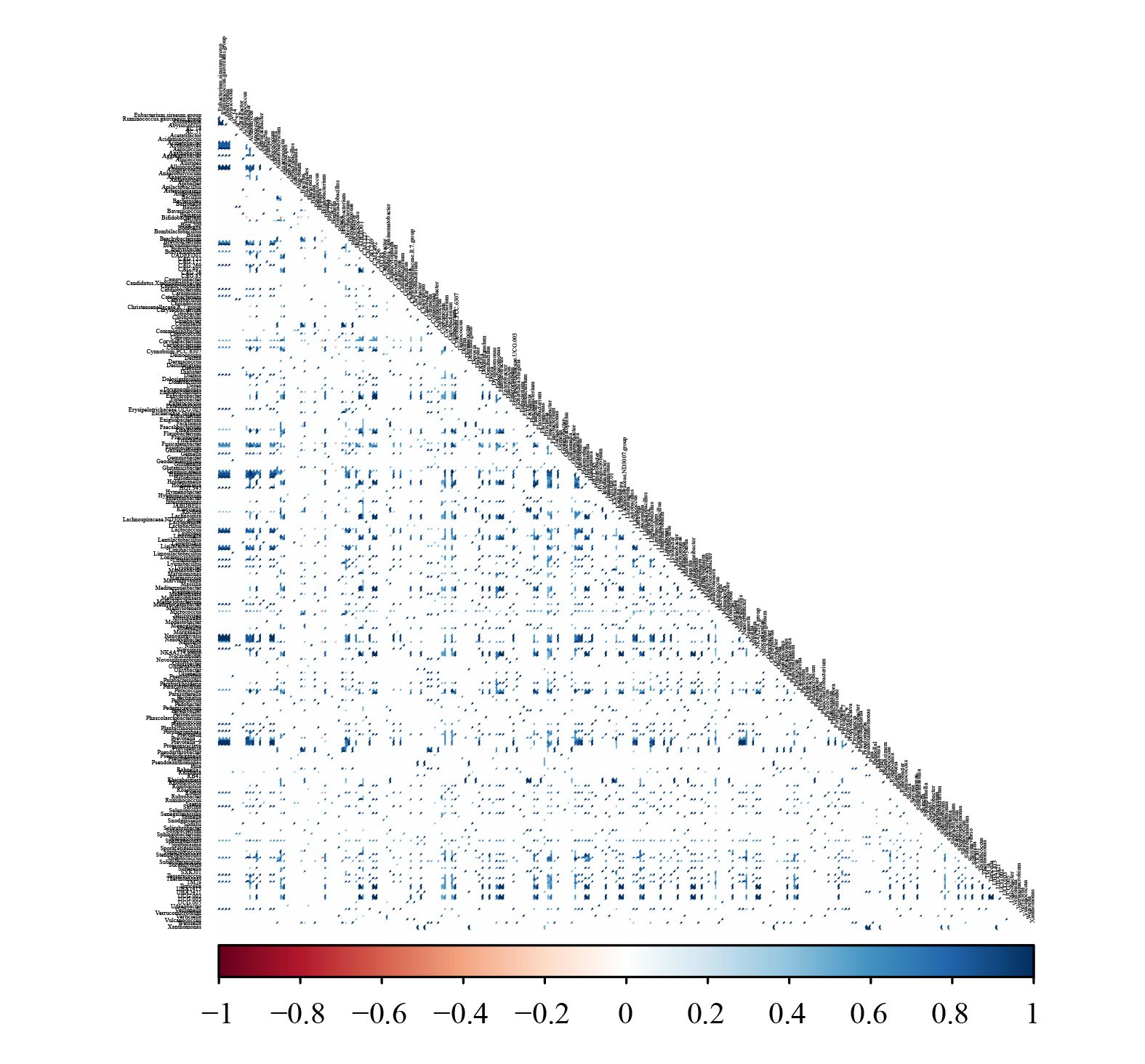

**Fig S5.** **Co-occurrence of bacterial genera across all the honey bee species**. Axis represents microbial taxa and each point indicates pairwise co-occurrence strength between taxa (Blue colour denotes positive correlation while red represents negative correlation.)

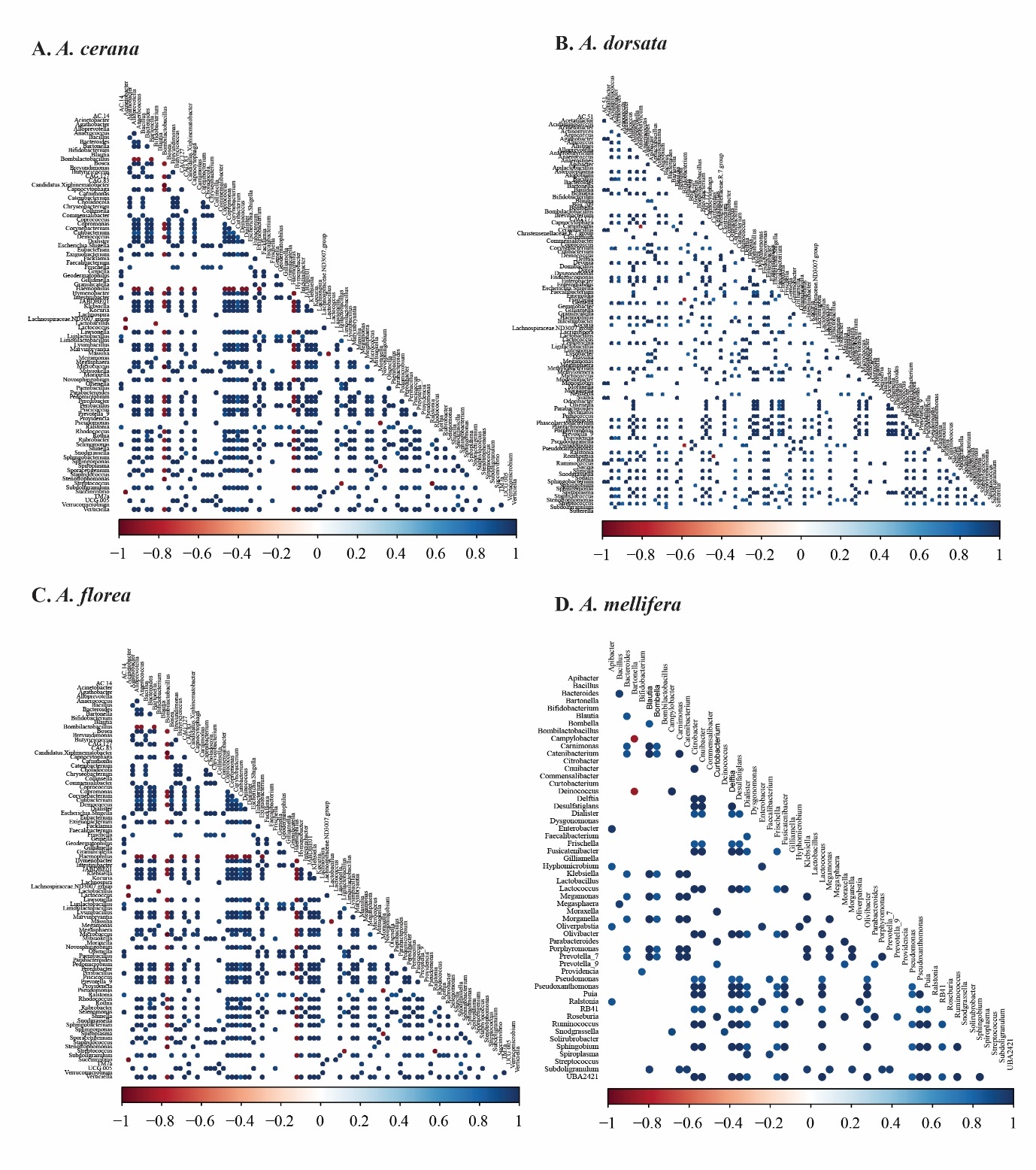
**Fig S6**. **Species-wise representation of cooccurrence of bacterial genera. (A)** *A. cerana***, (B)** *A. dorsata*, **(C)** *A. florea***, (D)** *A. mellifera.* Axis represents microbial taxa, and each point indicates pairwise co-occurrence strength between taxa (Blue colour denotes positive correlation, while red represents negative correlation) Dataset S5.

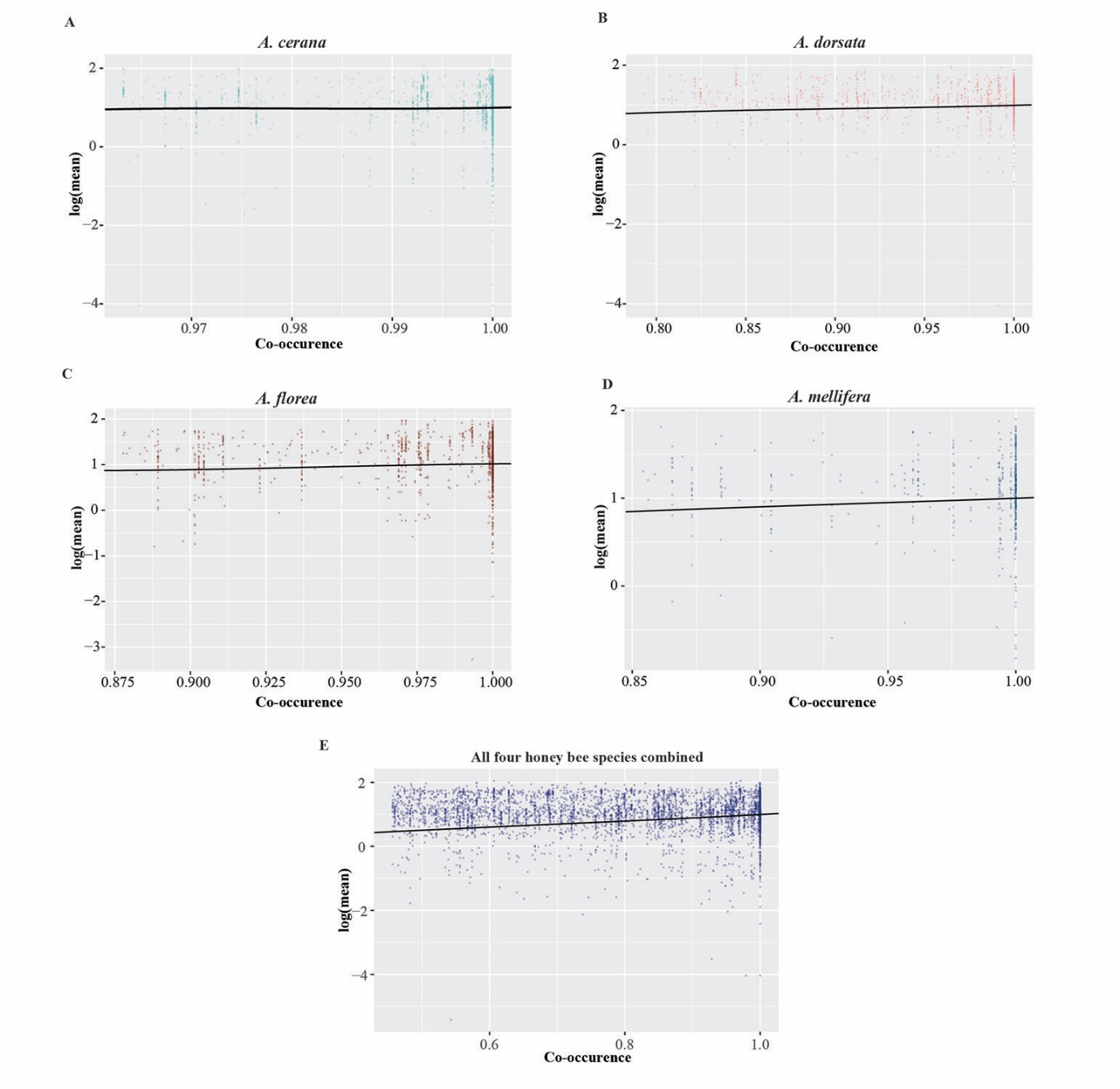

**Fig. S7. Association between phylogenetic relatedness and co-occurrence of bacterial genera across honey bee species.** (A) *A. cerana*, (B) *A. dorsata*, (C) *A. florea*, (D) *A. mellifera,* (E) All four species combined. Scatter plots show the relationship between pairwise co-occurrence and log-transformed phylogenetic distances between microbial taxa.

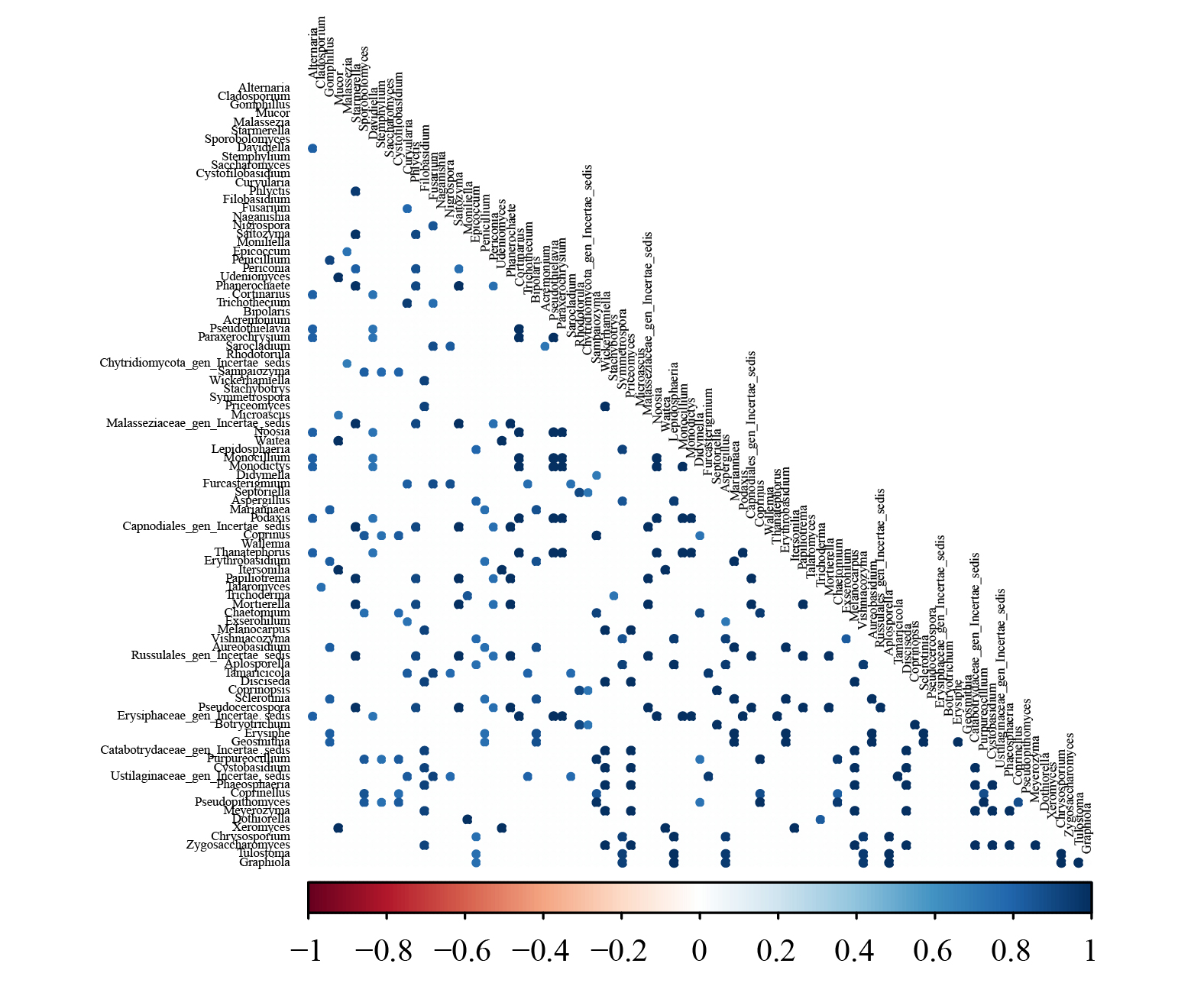

**Fig S8.** **Co-occurrence of fungal genera across honey bee species.** Axis represents microbial taxa and each point indicates pairwise co-occurrence strength between taxa (Blue colour denotes positive correlation while red represents negative correlation.)

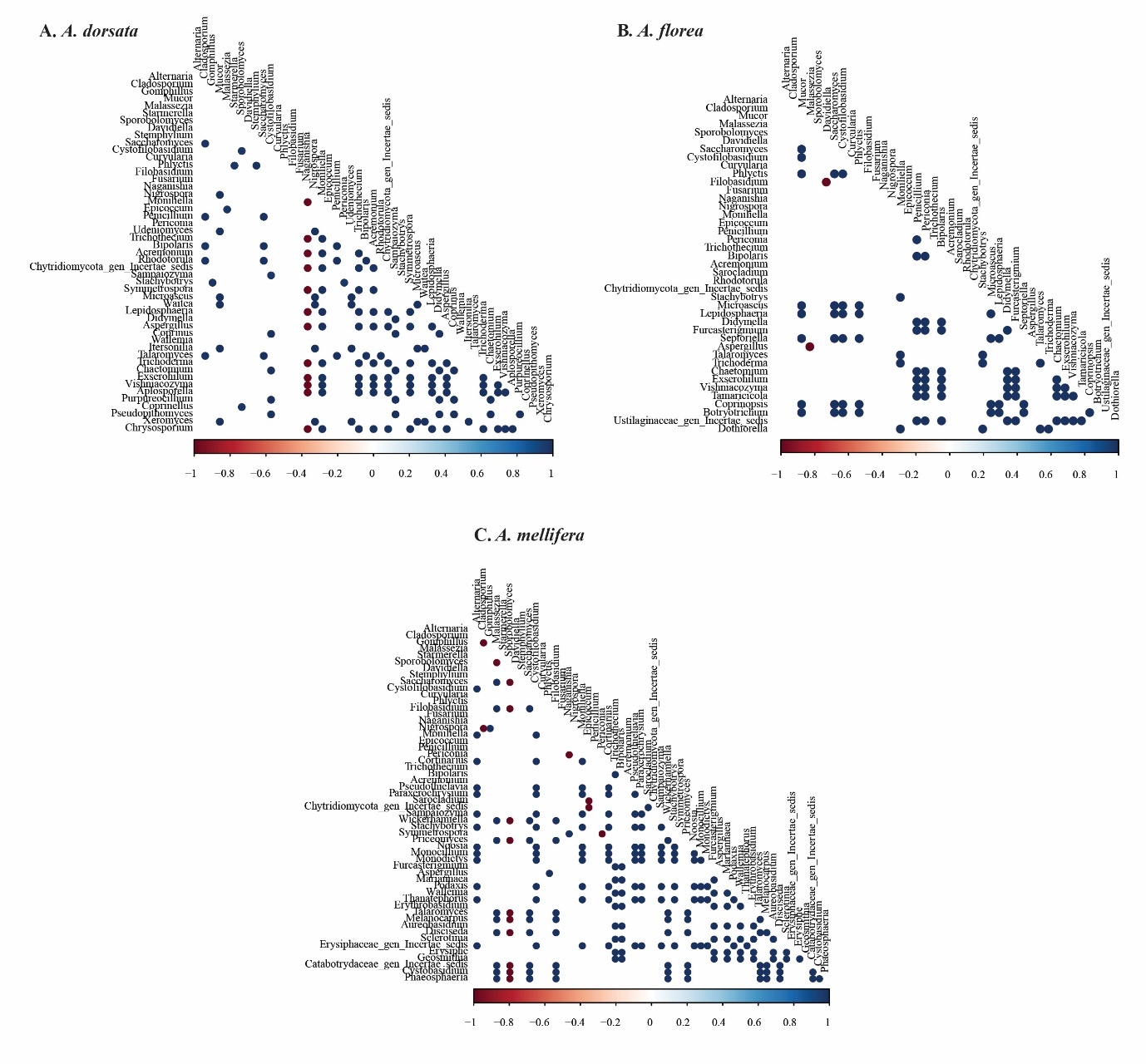
**Fig S9**. **Species-wise representation of co-occurrence of fungal genera. (A)** *A. dorsata,* **(B)** *A. florea***, (C)** *A. mellifera.* Axis represents microbial taxa and each point indicates pairwise co-occurrence strength between taxa (Blue colour denotes positive correlation while red represents negative correlation) (Dataset S6).

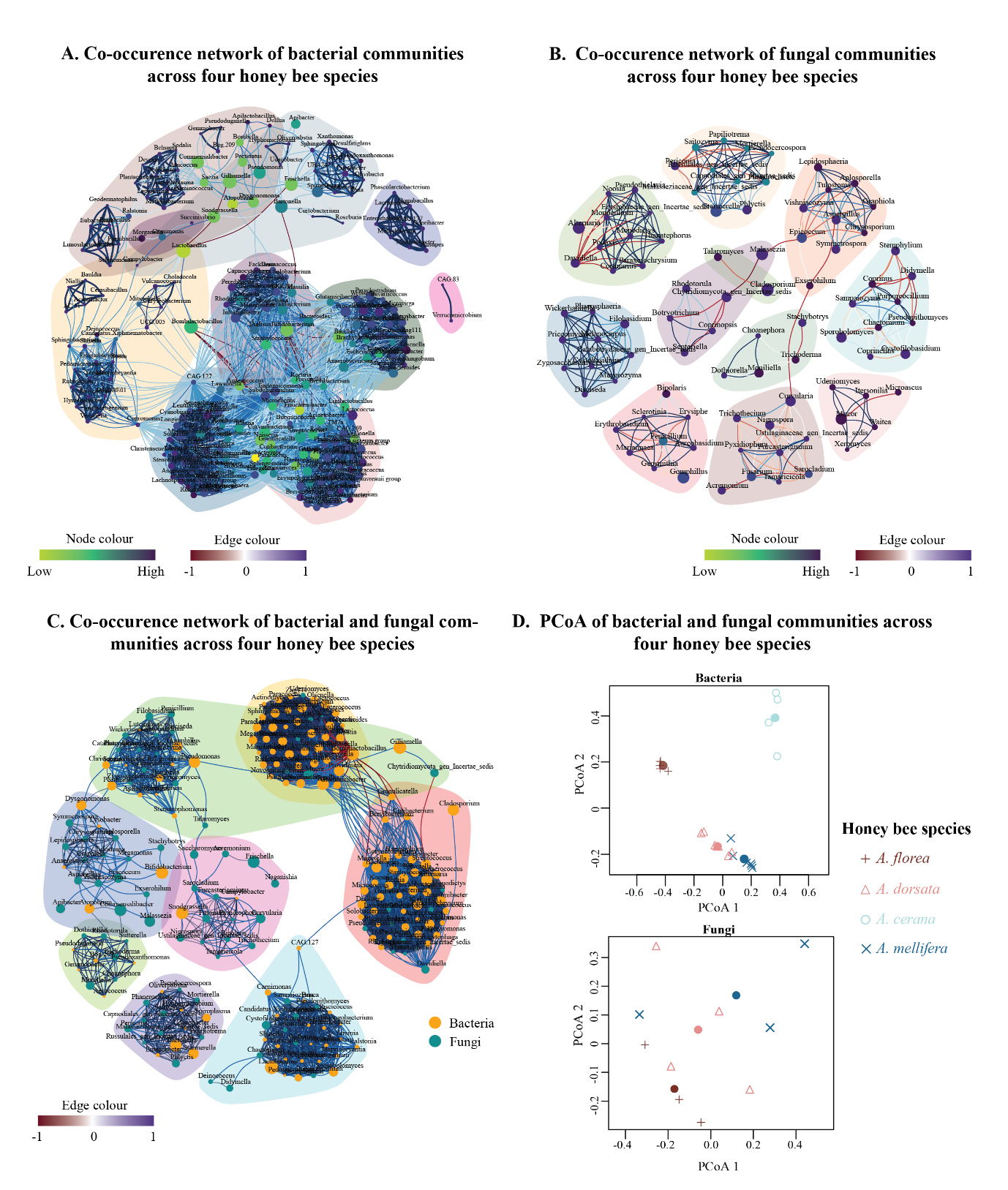

**Figure S10. Co-occurrence networks of microbial communities across four honey bee species.** (A) Bacteria (B) Fungi (C) Integrated bacterial–fungal network. In the (A) Bacteria and (B) Fungi co-occurrence networks, nodes represent microbial genera, with node colour ranging from cobiont homogeneity (yellow) to heterogeneity (blue), and node size proportional to relative abundance. In the integrated network (C), bacterial taxa are shown in orange, while fungal taxa are represented in green. Edges indicate co-occurrence relationships, with colour distinguishing positive and negative associations, and thickness reflecting the significance of correlations between genera. Shaded background regions denote distinct network modules. (D) Principal coordinate analysis (PCoA) of bacterial and fungal communities, based on the Bray-Kurtis analysis.

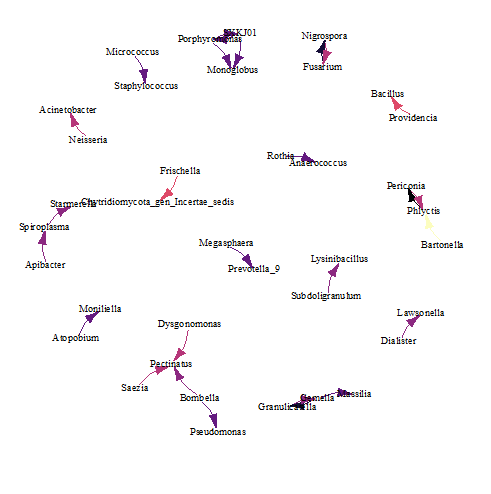

**Fig S11**. **Directional co-occurrence network of bacterial and fungal metacommunities inferred using a Bayesian network approach.** Arrows indicate probabilistic dependencies among taxa. Edge colour reflects the strength of the inferred relationships, with higher probability of directional association shown in yellow and lesser probabilistic associations in blue.

**SUPPLEMENTARY DATASETS**

**Dataset S1.** Metadata file containing the species information, Field IDs, SRA IDs, Bio accession number, read counts, and percentage of reads that were used for assigning the taxonomy.

**Dataset S2.** Mean abundance and average foraging time of bee species across all field sites.

**Dataset S3**. Relative abundance profiles and richness at the genus level of bacterial and fungal communities across field sites.

**Dataset S4**. Functional pathway abundance from PICRUSt and comparison across bees.

**Dataset S5.** Co-occurrence network of bacterial communities inferred from all four honeybee species.

**Dataset S6.** Co-occurrence network of fungal communities inferred from three honeybee species.
